# Endoplasmic reticulum associated degradation preserves hematopoietic stem cell quiescence and self-renewal by restricting mTOR activity

**DOI:** 10.1101/709964

**Authors:** Lu Liu, Ayaka Inoki, Kelly Fan, Fengbiao Mao, Guojun Shi, Xi Jin, Meiling Zhao, Gina Ney, Shengyi Sun, Yali Dou, Ken Inoki, Ling Qi, Qing Li

**Author notes:** Corresponding Author and lead contact: Qing Li.

## Abstract

Many tissue-specific stem cells require quiescence to sustain stem cell pool and maintain lifelong tissue integrity. It remains unclear whether protein quality control is required for stem cells in quiescence when RNA content, protein synthesis and metabolic activities are significantly reduced. Here, we report that endoplasmic reticulum associated degradation (ERAD) is required to preserve the function of quiescent hematopoietic stem cells (HSC). The Sel1L/Hrd1 ERAD genes are enriched in the quiescent and inactive HSCs, and conditional knockout of Sel1L in hematopoietic tissues drives HSCs to hyper-proliferation which leads to reduced self-renewal and HSC depletion. ERAD deficiency induces a non-apoptotic ER stress and activates unfolded protein response (UPR). Furthermore, Sel1L knockout leads to mTOR activation, and mTOR inhibition rescues the HSC defects in Sel1L knockout mice. Protein quality control is, therefore, tightly regulated and actively engaged in quiescent HSCs. Sel1L/Hrd1 ERAD maintains HSC quiescence and self-renewal via restricting mTOR activity.

## Introduction

Most adult tissue-specific stem cells persist in a quiescent state for a prolonged period of time (Li and Clevers, 2010; van Velthoven and Rando, 2019), and loss of stem cell quiescence can lead to impaired stem cell functions. For instance, loss of quiescence in hematopoietic (Cheng et al., 2000; Ito et al., 2004; Karlsson et al., 2007; Kharas et al., 2010; Maryanovich et al., 2012; Matsumoto et al., 2011; Wilson et al., 2004; Yilmaz et al., 2006; Zhang et al., 2006), muscle (Bjornson et al., 2012; Cheung et al., 2012; Mourikis et al., 2012), or neural stem cells (Bjornson et al., 2012; Farioli-Vecchioli et al., 2012; Jones et al., 2015), leads to reduced stem cell self-renewal, depletion of the stem cell pool and failure in tissue regeneration. Upon entering quiescence, stem cells reduce the cell size, become less transcriptionally and metabolically active, and exhibit reduced RNA contents and lower protein synthesis rate (Buszczak et al., 2014; Llorens-Bobadilla et al., 2015; Signer et al., 2014; Zismanov et al., 2016). This raises a major question whether protein quality control is actively involved in maintaining stem cell functions when stem cells are in the deeply dormant state.

Endoplasmic Reticulum has emerged as a central site where protein homeostasis is maintained by integrating protein synthesis, folding/processing, trafficking and degradation control systems. Perturbation in protein homeostasis in ER leads to increased ER stress with higher concentration of unfolded or misfolded proteins, which activates a cascade of signal transduction pathways termed unfolded protein response (UPR) (Hetz, 2012; Ron and Walter, 2007). Three main branches of signal transduction have been identified in mammalian cells in response to ER stress: inositol-requiring enzyme 1α (IRE1α, gene is also known as *ERN1*), pancreatic eIF-2α kinase (PERK) and activating transcription factor 6 (ATF6). Activation of UPR either restores the balance of protein homeostasis, by constraining protein synthesis and increasing protein folding and degradation, or triggers apoptosis if a cell is deemed beyond repair after prolonged stress. The cell context and the intensity and duration of the cellular stress determine the fate of the cell following ER stress (Hetz, 2012; Ron and Walter, 2007).

Another important mechanism of ER stress response, and proposed downstream target of UPR, is the ER associated degradation (ERAD) system. These ER transmembrane protein complexes recognize misfolded proteins in ER and dislocate them to cytosolic compartment to undergo ubiquitination for proteasome degradation (Hwang and Qi, 2018; Ruggiano et al., 2014). There is a constant demand for ERAD-mediated surveillance to clear aberrant proteins in the ER, which occur when a native structure fails to achieve due to mutation, translational mis-incorporation or stochastic inefficiency in adopting native conformation or forming protein complexes. Sel1L, is an important adaptor of the Hrd1 ERAD complex that recognizes misfolded proteins in ER lumen and recruits them to ERAD complexes for dislocation to cytosol for proteasome degradation. Sel1L has also been shown to bind to and stabilize Hrd1, an E3 ligase of the ERAD complex, therefore it’s indispensable for Hrd1 ERAD function (Qi et al., 2017). The function of Sel1L/Hrd1 ERAD has been studied in many differentiated cell types including liver, adipocytes and B cells (Bhattacharya et al., 2018; Francisco et al., 2010; Ji et al., 2016; Sha et al., 2014; Sun et al., 2016; Sun et al., 2014; Sun et al., 2015). The function of ERAD in tissue stem cells has not been well studied.

Hematopoietic stem cells (HSCs) are a rare population of cells that reside in the bone marrow niche and are at the apex of hematopoietic hierarchy (Mendelson and Frenette, 2014; Morrison and Scadden, 2014; Orkin and Zon, 2008). The highest long-term self-renewal potential is preserved in the most quiescent HSCs (Foudi et al., 2009; Wilson et al., 2008; Wilson et al., 2009). Similarly, HSCs that display higher level of mitochondrial activity give rise to much lower levels of blood cell reconstitution in transplant recipients than HSCs with low mitochondrial activity (Vannini et al., 2016), indicating that HSC self-renewal potential is retained in metabolically inactive subpopulation. Furthermore, signals that drive HSCs into proliferation cycle often lead to HSC differentiation and exhaustion (Cheng et al., 2000; Ito et al., 2004; Karlsson et al., 2007; Kharas et al., 2010; Maryanovich et al., 2012; Matsumoto et al., 2011; Wilson et al., 2004; Yilmaz et al., 2006; Zhang et al., 2006). Imbalance of HSC quiescence, proliferation and differentiation can lead to hematopoietic failure or malignancies (Reya et al., 2001; Rossi et al., 2008). Although the majority of HSCs are quiescent and exhibit extremely low level of protein synthesis and therefore low burden for protein folding, our group and others recently reported that UPR pathways, such as IRE1α/XBP1s and PERK/eIF2α are important regulators of HSC functions under stress conditions when protein homeostasis is perturbed (Liu et al., 2019; van Galen et al., 2014; van Galen et al., 2018). At steady state, less than 2% of long-term HSCs are actively cycling (in S-G2-M phase of cell cycle) (Foudi et al., 2009; Wilson et al., 2008; Wilson et al., 2009), raising a major question whether protein quality control is important for HSCs at steady state under physiological conditions. Furthermore, within the overall quiescent long-term HSC pool, a fraction of cells (∼20% based on label retention assay) divides at an extremely low rate (Foudi et al., 2009; Wilson et al., 2008; Wilson et al., 2009). It remains unclear if protein quality control is required for HSCs that are in the deeply dormant state.

Here we report that the protein quality control system Sel1L/Hrd1 ERAD is actively engaged in quiescent HSCs to preserve HSC self-renewal. Sel1L, and other members of the ERAD complex, are consistently enriched in quiescent and metabolically inactive HSCs. Deletion of Sel1L leads to HSC proliferation and activation, resulting in loss of HSC reconstitution and the depletion of the HSC pool. Loss of Sel1L induces non-apoptotic ER stress and activates UPR signaling. Activation of IRE1α/XBP1 and PERK/eIF2α branches of UPR preserves the function of Sel1L knockout HSCs, indicating these branches are compensatory pathways to protect Sel1L KO HSCs under ER stress. On the other hand, Sel1L KO leads to AKT-mTOR activation, and inhibition of mTOR signaling restores the functions of Sel1L knockout HSCs in vivo. Our studies identify Sel1L/Hrd1 ERAD as a critical protein quality control pathway that maintains HSCs in quiescence and determines HSC fate by modulating mTOR activity.

## Results

### The Sel1L-Hrd1 ERAD genes are enriched in dormant and metabolically inactive HSCs

To determine the role of protein quality control in HSC quiescence, we examined the protein aggregates in quiescent/dormant (dHSC) versus proliferative/activated (aHSC) HSCs in the *Col1α1-H2B-GFP; Rosa26-M2-rtTA* double transgenic mice (**Figure S1a** and Figure 1a) (Foudi et al., 2009). In this model, HSCs were first labeled with Histone 2B-green fluorescent protein (H2B-GFP) during a 6-week period of doxycycline administration. After removal of doxycycline, HSCs dilute GFP with each round of division, which allows tracking of proliferation over time. The aHSCs (GFP low; GFP^lo^) and dHSCs (GFP high; GFP^hi^) can then be separated by GFP expression after a 12-18 week ‘off labeling’ chase period (**Figure S1a** and Figure 1a). First, we observed that protein aggresome level, as measured by PROTEOSTAT (Enzo life sciences), was much higher in aHSCs (GFP^low^) than dHSCs (GFP^high^), suggesting that protein homeostasis is tightly controlled in quiescent HSCs (Figure 1a-b). To determine whether protein quality control pathways play any role in regulating HSC quiescence, we examined the published gene expression profile comparing dHSCs and aHSCs (Cabezas-Wallscheid et al., 2017). While proteasome degradation pathways are highly active in aHSCs as reported (Cabezas-Wallscheid et al., 2017), the levels of many ERAD genes, such as Herpud1, Hrd1 and Sel1L, are enriched in dHSCs (**Figure S1b**). In addition, our previously published gene expression data (Li et al., 2013a)(GSE45194) revealed that the level of Sel1L is much higher in quiescent HSCs (Figure 1c), suggesting an important role of ERAD in regulating HSC quiescence. The increased expression of Sel1L in dHSCs was confirmed by quantitative RT-qPCR on purified dHSC (GFP^high^) as compared to aHSCs (GFP^low^) (Figure 1d). When HSCs exit quiescence and become metabolically activated, they exhibit higher levels of mitochondrial membrane potential, which can be measured by tetramethyl rhodamine methyl ester (TMRM) (**Figure S1c-d**)(Vannini et al., 2016). Next, we purified metabolically inactive (TMRM^lo^) versus active (TMRM^hi^) SLAM HSCs, and compared the expression of ERAD genes (Figure 1e). Sel1L level was significantly reduced in metabolically active HSCs (Figure 1f) and was further decreased in progenitors (**Figure S1e)**. Taken together, Sel1L/Hrd1 genes are enriched in quiescent and metabolically inactive HSCs.

**Figure 1.**
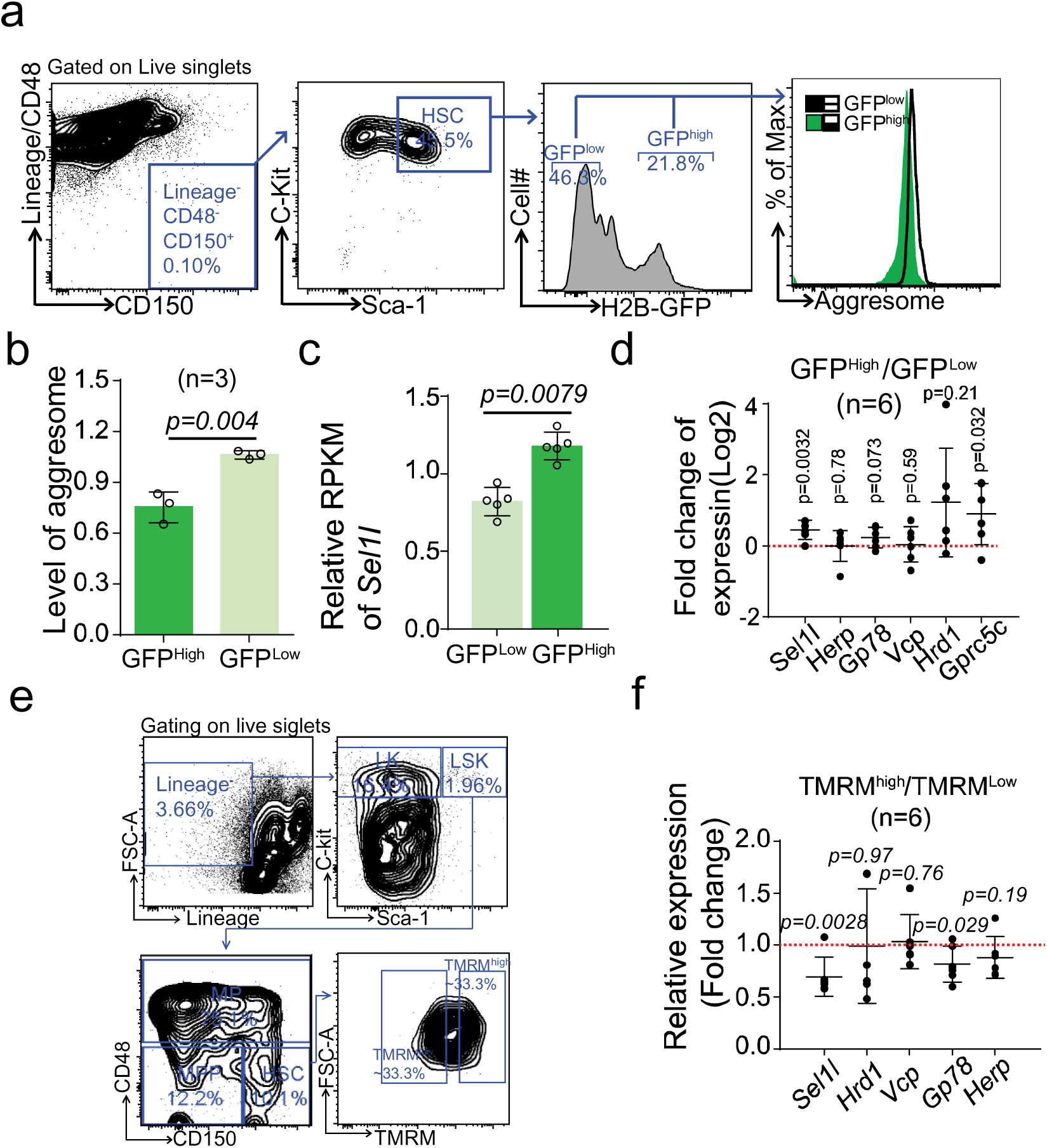
Sel1l/Hrd1 ERAD genes are enriched in quiescent and dormant HSCs. **a-b**, *Col1α1-H2B-GFP^+/-^; Rosa26-M2-rtTA^+/-^* mice were labeled with doxycycline for 6 weeks and then “off label chase” for 18 weeks. H2B-GFP label retention allows purification of quiescent (GFP^high^) and proliferative (GFP^low^) HSCs. Aggresome levels in different sub-populations were detected by PROTESTAT staining. **a**, Representative FACS plots gating strategy for sorting and analyzing. **b**, Summary of protein aggregation level in quiescent (GFP^high^) and proliferative (GFP^low^) HSCs relative to bulk HSCs. Data represent mean±s.d. **c**, Sel1L level from Microarray analysis comparing quiescent (GFP^high^) and proliferative (GFP^lo^) HSCs (Li et al., 2013b). Data represent Reads Per Kilobase of transcript per Million mapped reads (RPKM), each point represents a single probe. *p* value determined by two-sided Wilcoxon rank sum test. **d**, qRT-PCR of ERAD genes in quiescent (GFP^hi^) versus proliferating (GFP^low^) HSCs purified from *Col1α1-H2B-GFP^+/+^; Rosa26-M2-rtTA^+/+^*. Data represent mean±s.d. Each point represents fold change of FACS-sorted quiescent HSCs (GFP^high^) versus proliferative HSCs (GFP^low^) from a single mouse. **e-f**, metabolically inactive and active sub-populations of HSCs in 8-10 weeks old wildtype mice were purified based on tetramethyl rhodamine methyl ester (TMRM) staining. **e,** Representative FACS plots and gating strategy for sorting. **f,** qRT-PCR of ERAD genes in metabolically active (TMRM^high^) versus metabolically inactive (TMRM^low^) HSCs. Data represent mean±s.d. Each point represents fold change of FACS-sorted metabolic active HSCs (TMRM^high^) versus metabolic quiescent (TMRM^low^) HSCs from a single mouse. Two-sided student t-test was used for statistical analysis unless specified.

### Sel1L-Hrd1 ERAD deficiency depletes HSC pool but expands lineage restricted progenitors

To determine the role of Sel1L/Hrd1 ERAD in hematopoiesis, we crossed *Mx1-Cre^+^* mice with *Sel1L^fl/fl^* mice (Sun et al., 2014), and injected 6-8 weeks old *Mx1-cre^+^; Sel1L^fl/fl^* mice with poly-inosine-poly-cytosine (pIpC; 0.5 ug/gram body weight, every other day for 6 doses) to delete Sel1L in hematopoietic tissues. Sel1L knockout (KO) was confirmed by Western Blot on whole bone marrow cells or lineage^-^Sca1^+^cKit^+^ (LSK) cells from *Mx1-Cre^+^; Sel1L^fl/fl^* mice (**Figure S2a** and Figure 7a). Sel1L KO mice exhibited similar levels of body weight, bone marrow and spleen cellularity, but slightly increased spleen weight and a significant reduction of thymus weight and cellularity (Figure 2a-f and **Figure S2b-d**). The thymic defect is consistent with a previous study reporting that loss of Hrd1 leads to T cell developmental defects (Xu et al., 2016). We next enumerated the hematopoietic populations in the bone marrow and spleen of Sel1L KO mice by FACS analysis. We observed significantly reduced numbers and frequencies of SLAM HSCs (Kiel et al., 2005b) (CD150^-^CD48^-^LSK) and multi-potent progenitors (MPP; CD150^+^CD48^-^LSK) in the bone marrow (Figure 2g-h and **Figure S2e**). The numbers and frequencies of broader progenitor population as measured by LSK were not affected in Sel1L knockout bone marrow but were increased in the spleen (Figure 2g-h and **Figure S2e-f**). The myeloid progenitor granulocyte-macrophage progenitor (GMP; Lineage^-^Sca1^-^cKit^+^ CD16/32^+^CD34^+^) and the common lymphoid progenitor population (CLP; Lineage^-^Sca1^+^cKit^+^ IL7Ra^+^FLT3^+^) were expanded in Sel1L KO bone marrow, while all progenitor populations including common myeloid progenitor (CMP; Lineage^-^Sca1^-^cKit^+^ CD16/32^+^CD34^-^), megakaryocyte-erythroid progenitor (MEP; Lineage^-^Sca1^-^cKit^+^ CD16/32^-^CD34^-^), GMP and CLP were expanded in the spleen of Sel1L KO mice (Figure 2i-j and **Figure S2g-h**). These findings suggest that Sel1L is required to maintain HSC pool and loss of Sel1L leads to depletion of HSC and MPPs and expansion of lineage restricted progenitors. Analysis of mature myeloid, B and T cells revealed reduced frequency of B cells in the spleen and bone marrow (Figure 2k-l and **Figure S2i-k**), which is consistent with a recent report (Ji et al., 2016), and no significant change of the frequency of myeloid and T cells. Consistently, Vav1-cre mediated Sel1L deletion lead to similar HSC depletion and altered hematopoiesis (Figure 3 **and Figure S3**). However, significantly reduced body weight and bone marrow cellularity were observed in the *Vav1-cre^+^; Sel1L^fl/fl^* mice by 6-8 weeks of age (Figure 3a, 3d, **Figure S3a, S3d**). We opted to focus further analysis of HSC and progenitors on the *Mx1-cre^+^; Sel1l^fl/fl^* mice.

**Figure 2.**
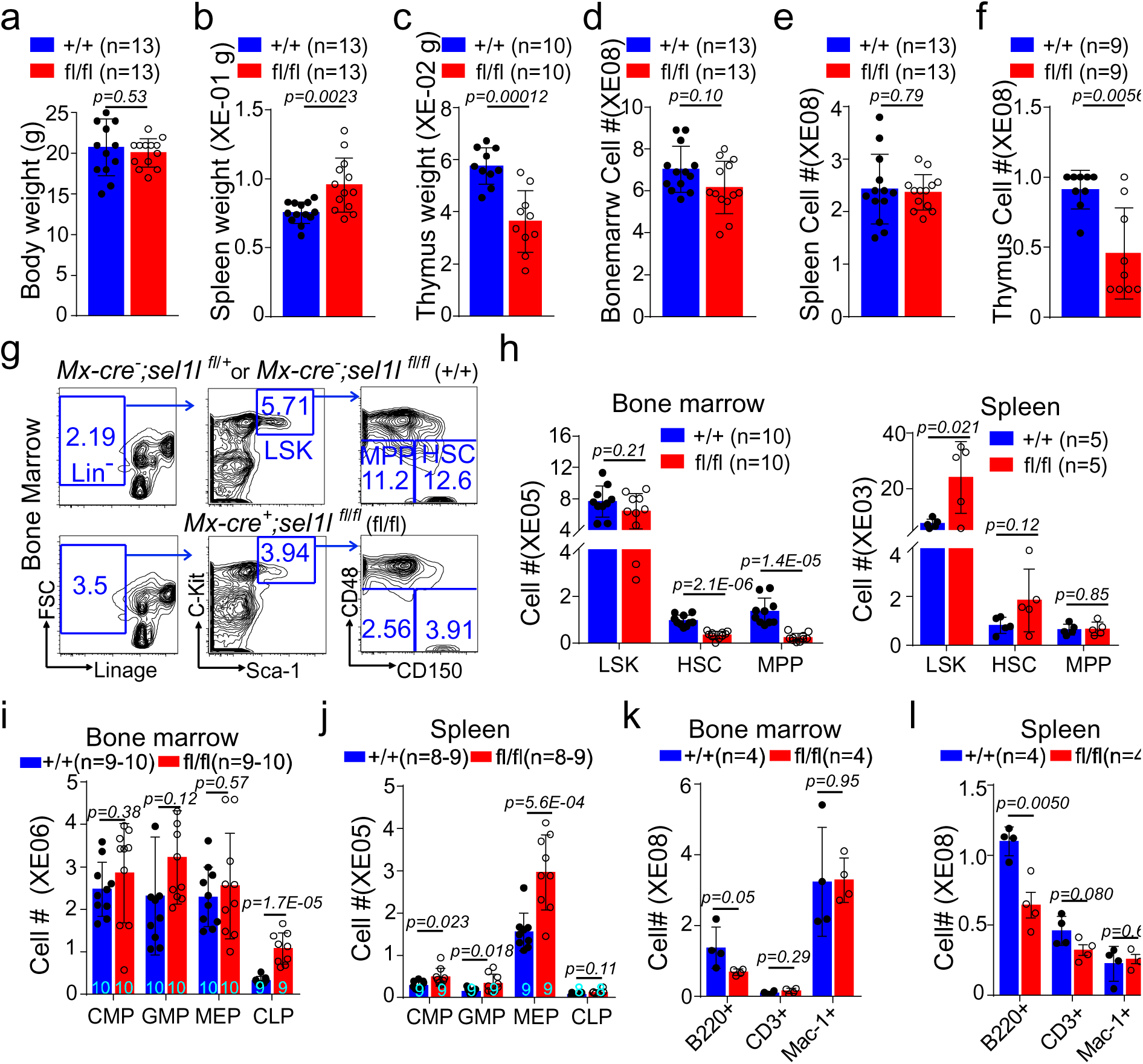
Sel1l-Hrd1 ERAD is required to maintain HSC pool. 6-8 weeks old *Mx1-cre^-^; Sel1L^fl/fl^* or *Mx1-cre^-^; Sel1L^fl/+^* (+/+) and *Mx1-cre^-^; Sel1L^fl/fl^* (fl/fl) mice were injected with pIpC every other day for a total of 6 doses. Two weeks after pIpC injection, body weight **(a)**, spleen and thymus weight (**b, c)**, cellularity of bone marrow, spleen and thymus (**d-f**), numbers of HSPC (**g-h**), lineage restricted progenitors (**i, j**) and mature blood cells (**k, l**) were analyzed. Data represent mean±s.d. Each replicate represents a single mouse from an independent experiment. Two-sided student t-test was used for statistical analysis unless specified.

**Figure 3.**
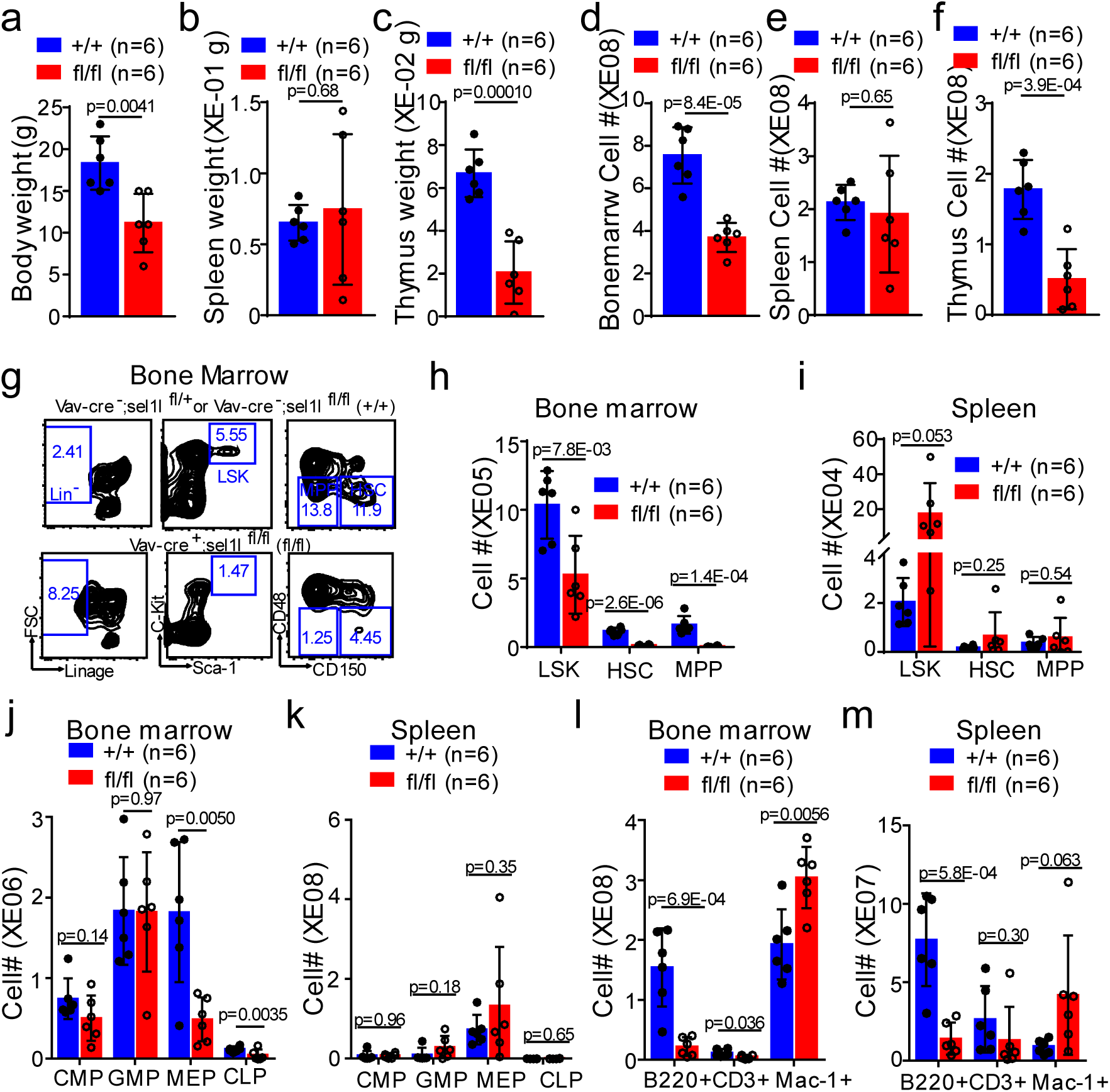
Steady state hematopoiesis in vav-cre+; Sel1lfl/fl mice. 6 to 8 weeks old *Vav-cre+; Sel1l^fl/fl^ (fl/fl)* and *Vav-cre^-^; Sel1l^fl/+^* or *Vav-cre^-^; Sel1l^fl/fl^* (+/+) mice were analyzed for body weight (**a**), spleen and thymus weight (**b, c**), cellularity of bone marrow, spleen and thymus (**d-f**), numbers of HSPC (**g-i**), lineage restricted progenitors (**j-k**) and mature blood cells (**l-m**). Data represent mean±s.d. Each replicate represents a single mouse from an independent experiment. Two-sided student t-test was used for statistical analysis unless specified.

### Sel1L-Hrd1 ERAD deficiency leads to reduced HSC reconstitution

To determine the role of Sel1L in HSC reconstitution and self-renewal, we performed competitive repopulation assay. Two weeks after pIpC injection, we transplanted 3×10^5^ whole bone marrow cells from *Mx1-cre^+^; Sel1L^fl/fl^* or control mice (CD45.2^+^), together with 3×10^5^ CD45.1^+^ wild type bone marrow cells, into lethally irradiated CD45.1^+^ wild-type recipients. Donor reconstitution in total CD45+, myeloid (Mac1^+^), B (B220^+^) or T (CD3^+^) cells in the peripheral blood was analyzed every 4 weeks for 16 weeks after transplantation (**Figure S4a**). Sel1L deficiency resulted in significantly lower levels of long-term multi-lineage donor reconstitution in transplant recipients (Figure 4a). To confirm the reduction of HSC reconstitution and self-renewal, the levels of donor HSCs, progenitors, and lineage cells in the bone marrow were analyzed in transplant recipients at the end of 16 weeks. This showed that all donor populations were significantly reduced in Sel1L KO transplant recipients compared to control mice (Figure 4b). Because the overall lower HSC reconstitution in transplant recipients in the above experiment may be attributed to the lower HSC frequency in the bone marrow of Sel1L KO mice. To assess the function of individual HSCs, we transplanted 50 FACS-purified HSCs from either Sel1L KO or control mice into irradiated receipt mice along with 0.2 million CD45.1 whole bone marrow competitor cells (**Figure S4b)**. Sel1L KO HSCs failed to yield any long-term reconstitution in transplant recipients (Figure 4c). Again, consistent with reduced HSC reconstitution and self-renewal, the levels of donor HSCs, progenitors, and lineage cells in the bone marrow of Sel1L KO transplant recipients were significantly reduced compared to controls (Figure 4d).

**Figure 4.**
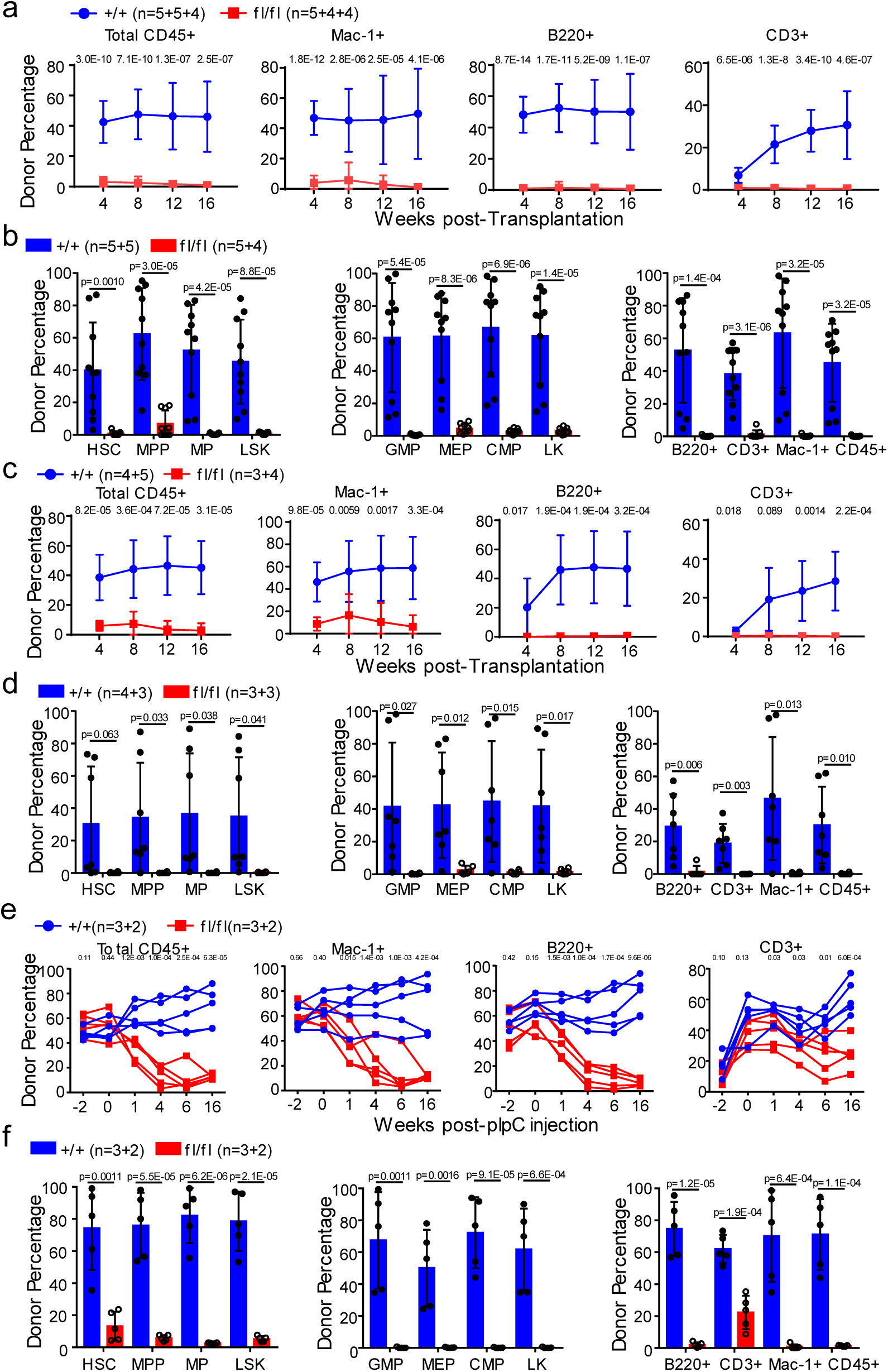
Sel1L-Hrd1 ERAD is required to preserve HSCs functions. **a**, whole bone marrow cells (0.3 million) from CD45.2 *Mx1-cre^+^; Sel1L^fl/fl^* (fl/fl) or control (+/+) mice were transplanted into lethally irradiated CD45.1 mice together with 0.3 million CD45.1 competitor cells. The contribution of CD45.2 cells in total CD45+, myeloid (Mac-1^+^), B (B220^+^) and T (CD3^+^) cells was analyzed from peripheral blood. **b**, Donor contribution to HSCs and other hematopoietic populations in the bone marrow was analyzed in transplant recipients of *Mx1-cre^+^; Sel1L^fl/fl^* (fl/fl) or control (+/+) cells 16 weeks after transplantation. **c**, competitive repopulation assay with 50 FACS-purified HSCs from CD45.2 *Mx1-cre^+^; Sel1L^fl/fl^* (fl/fl) or control (+/+) mice transplanted into irradiated receipt mice along with 0.2 million CD45.1 whole bone marrow competitors. The contribution of CD45.2 cells in total CD45+, myeloid (Mac-1^+^), B (B220^+^) and T (CD3^+^) cells was analyzed from peripheral blood. **d**, Donor contribution to HSCs and other hematopoietic populations was analyzed in the bone marrow of transplant recipients of *Mx1-cre^+^; Sel1L^fl/fl^* (fl/fl) or control (+/+) HSCs 16 weeks after transplantation. **e**, chimerism maintenance analysis with 5×10^5^ CD45.2^+^ whole bone marrow cells from *Mx1-cre^+^; Sel1L^fl/fl^* or control mice (without pIpC) transplanted together with 5×10^5^ CD45.1^+^ wild type bone marrow cells into lethally irradiated CD45.1^+^ wild-type recipients. Transplants were injected with pIpC (3 doses) 6 weeks after transplantation. The contribution of CD45.2 cells in total CD45+, myeloid (Mac-1^+^), B (B220^+^) and T (CD3^+^) cells was analyzed from peripheral blood. **f**, Donor contribution to HSCs and other hematopoietic populations was analyzed in the bone marrow of transplant recipients from **e**. Data represent mean±s.d. N is described as number of replicates from each round of transplant separated by the plus (+) sign. Two-sided student t-test was used for statistical analysis unless specified.

To determine whether Sel1L KO affects HSCs that have already established engraftment and to avoid confounding transplantation-related effects such as homing and engraftment defects, we performed chimerism maintenance analysis (**Figure S4c**). We transplanted 5×10^5^ CD45.2^+^ whole bone marrow cells from *Mx1-cre^+^; Sel1L^fl/fl^* or control mice (without pIpC injection), together with 5×10^5^ CD45.1^+^ wild type bone marrow cells, into lethally irradiated CD45.1^+^ wild type recipients. Four weeks after transplantation, donor cells from all genotypes gave rise to similar levels of donor contribution (chimerism) to blood cells (Figure 4e). The recipients were then injected with pIpC (0.5 ug/gram body weight, every other day for 3 doses) 6 weeks after transplantation to knockout Sel1L. This led to a significant and rapid reduction of Sel1L KO donor chimerism in blood cells within 4-6 weeks after pIpC injection (Figure 4e), suggesting that Sel1L-Hrd1 ERAD is indispensable for HSCs reconstitution potential. We also observed significant reduction of donor HSCs, progenitors, and lineage cells in the bone marrow in Sel1L KO transplants (Figure 4f).

### Loss of Sel1L drives HSC to enter proliferation cycle and become metabolically activated

Our data showed that the expression level of Sel1L is much higher in quiescent HSCs as compared to proliferative or activated HSCs (Figure 1), suggesting a role of Sel1L in regulating HSC cycling. To evaluate the proliferative potential of Sel1L KO HSCs, we performed 5-bromo-2’-deoxyuridine (BrdU) incorporation assay. Following a 24-hour labeling with BrdU, Sel1L KO HSCs exhibited significantly higher frequency of BrdU^+^ HSCs (Figure 5a). To confirm the increased proliferation status of Sel1L KO HSCs, we performed H2B-GFP label retention assay by mating *Mx1-Cre^+^; Sel1L^fl/fl^* mice with *Col1A1-H2B–GFP; Rosa26-M2-rtTA* double transgenic mice (Figure 1a and **Figure S1a**) (Foudi et al., 2009). Two weeks after pIpC injection, Sel1L KO and control mice containing *Col1A1-H2B–GFP; Rosa26-M2-rtTA* double transgenes were placed on doxycycline water for 6 weeks, after which all HSCs were labeled with GFP (**Figure S1a**). Then doxycycline was removed to start “chase” period. After either a 12- or 18-week chase period, Sel1L KO HSCs retained much lower GFP levels compared to controls, indicating a higher overall proliferative state of the Sel1L KO HSCs (Figure 5b-c). In addition, mRNA levels of cell cycle inhibitors, p27 and p57, were reduced in Sel1L KO HSCs compared to control HSCs (**Figure S5a**). Taken together, loss of Sel1L/Hrd1 ERAD drives HSCs into hyper-proliferation.

**Figure 5.**
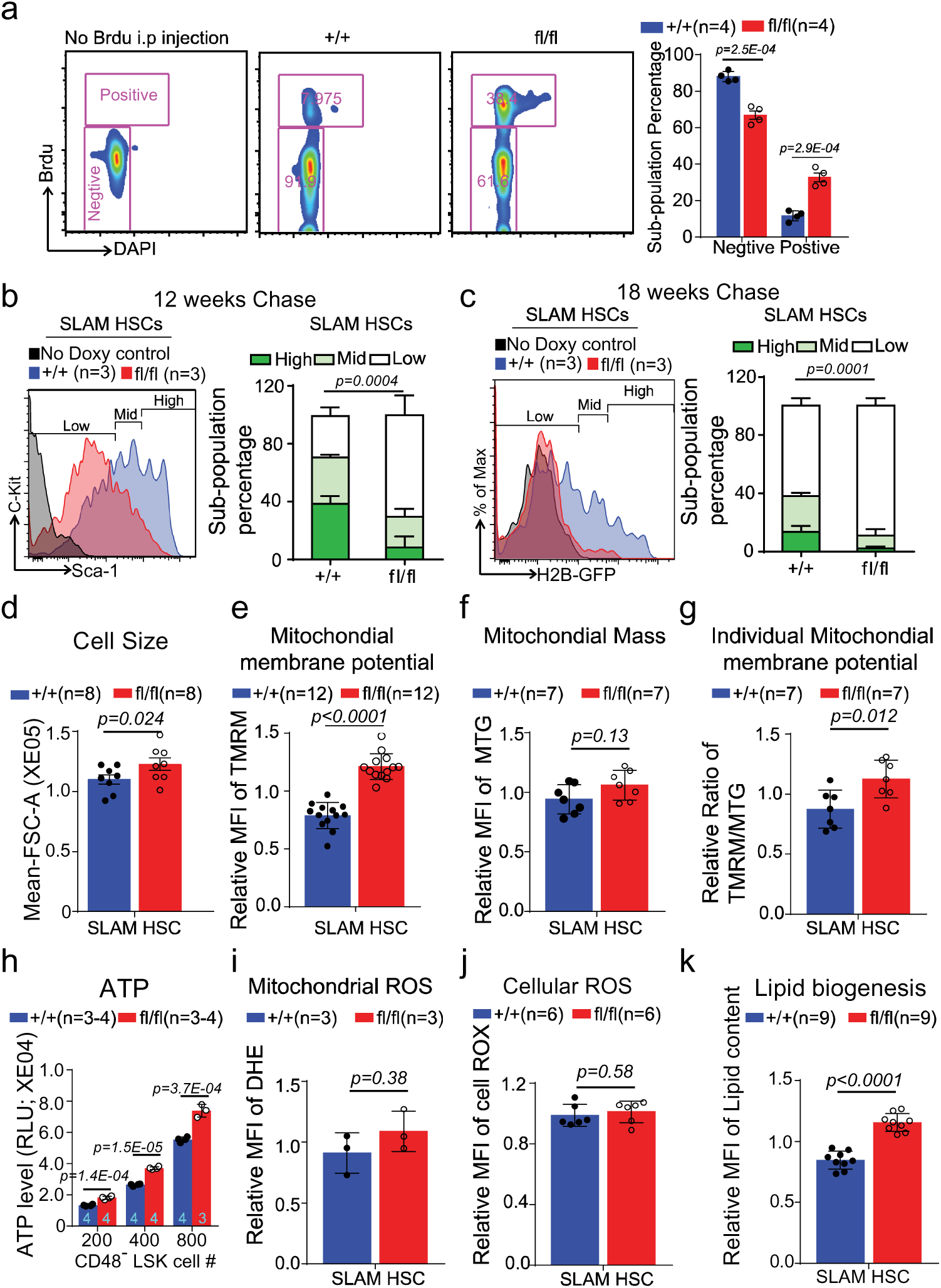
Loss of Sel1L leads to HSC hyper-proliferation and activation. **a**, Two weeks after pIpC injection, *Mx1-cre^+^; Sel1L^fl/fl^* (fl/fl) or control (+/+) mice were injected with BrdU (200mg/kg body mass; i.p.) and then placed on drinking water containing BrdU (1 mg/ml) for 24 hours. Representative FACS plots and gating strategy and summary of BrdU incorporation in HSCs are shown. **b-c**, Representative FACS plots with gating strategy, and summary of H2B-GFP retention assay. Two weeks after pIpC injection, *Mx1-cre^+^; Sel1L^fl/fl^*; *Col1A1-H2B–GFP^+/+^; Rosa26-M2-rtTA^+/+^* (fl/fl) or control (+/+) mice were place on doxycycline water (2g/L). After 6 weeks labeling, doxycycline water was removed and GFP level in HSCs was analyzed after 12 (**b**) or 18 (**c**) weeks “off-label” chase. **d-k**, Two weeks after pIpC injection, *Mx1-cre^+^; Sel1L^fl/fl^* (fl/fl) or control (+/+) mice were analyzed for cell size (by FSC-A, **d)**, mitochondrial membrane potential (by TMRM, **e**), mitochondrial mass (by Mito tacker green, **f**), ratio of mitochondrial membrane potential to mass (**g**), total ATP (by Cell Titer-Glo, **h**), mitochondrial ROS (by Dihydroergotamine mesylate, DHE, **i**), cellular ROS (by Cell Rox, **j**) and neutral lipid content (by HCS LipidTOX Neutral Lipid Stain, **k**). Data represent mean±s.d. Each point represents a single mouse from independent experiments. Two-sided student t-test was used for statistical analysis unless specified. For **e**-**g** and **i**-**k**, mean fluorescence intensity (MFI) were normalized to median value on the day of measurement, and *p* values were determined by two-sided Wilcoxon rank sum test.

Quiescent HSCs are more resistant than proliferative HSCs to repetitive proliferative stress, such as induced by 5-Fluorouracil (5FU) injection. To determine the effect of Sel1L knockout on the response of HSCs to proliferative stress, we injected *Mx1-Cre^+^; Sel1L^fl/fl^* and control mice with 5-FU at a dose of 150 mg/kg weekly (**Figure S5b**). Sel1L knockout mice demonstrated significantly shortened survival after 5-FU injection, consistent with a hyper-proliferative state and lower resistance to proliferative stress in Sel1L knockout HSCs.

Consistent with a higher proliferative and metabolically activated status, the Sel1L KO HSCs are significantly larger than control HSCs (Figure 5d), demonstrating increased mitochondrial membrane potential (measured by TMRM staining) and ATP content (measured by Cell Titer-Glo, Promega) (Figure 5e-h). Interestingly, the levels of mitochondrial and cellular ROS remain relatively unchanged in Sel1L KO HSCs (Figure 5i-j), suggesting a more balanced activation of redox metabolism. In addition, neutral lipid content was significantly higher in Sel1L KO HSCs, indicating increased lipid biogenesis (Figure 5k). Together with the reduced number of HSC and MPP and expanded downstream progenitor populations (Figure 2-3 and **S2-3**), these results indicate that ERAD machinery is required to maintain HSCs in quiescence, and ERAD deficiency, via Sel1L deletion, drives HSCs into proliferation and activation which leads to HSC differentiation.

### Sel1L-Hrd1 ERAD deficiency leads to non-apoptotic ER stress and activation of UPR

ERAD deficiency results in accumulation of misfolded ERAD substrate in ER to increase ER stress which activates unfolded protein response (UPR)(Hetz, 2012; Ron and Walter, 2007). To identify the mechanism through which Sel1L KO leads to HSC depletion, we determined if Sel1L KO leads to increased ER stress in HSCs. Sel1L KO HSCs exhibited increased level of protein aggresome (Figure 6a), and expansion of ER volume, consistent with increased ER stress (Figure 6b). To determine if Sel1L KO-induced ER stress leads to increased apoptosis, we measured the level of apoptosis in Sel1L KO HSCs by Annexin V staining. Sel1L KO and control HSCs exhibited similar level of apoptosis (Figure 6c), suggesting the loss of HSC function in Sel1L KO is unlikely due to increased apoptosis. In addition, Sel1L KO HSCs exhibited similar levels of apoptosis after treatment with ER stress inducers tunicamycin or thapsigargin (**Figure S6a-b**). These results demonstrate that, different from many ER stress conditions induced by extreme exogenous stress triggers, loss of Sel1L induces a non-apoptotic ER stress and represents a physiological stress.

**Figure 6.**
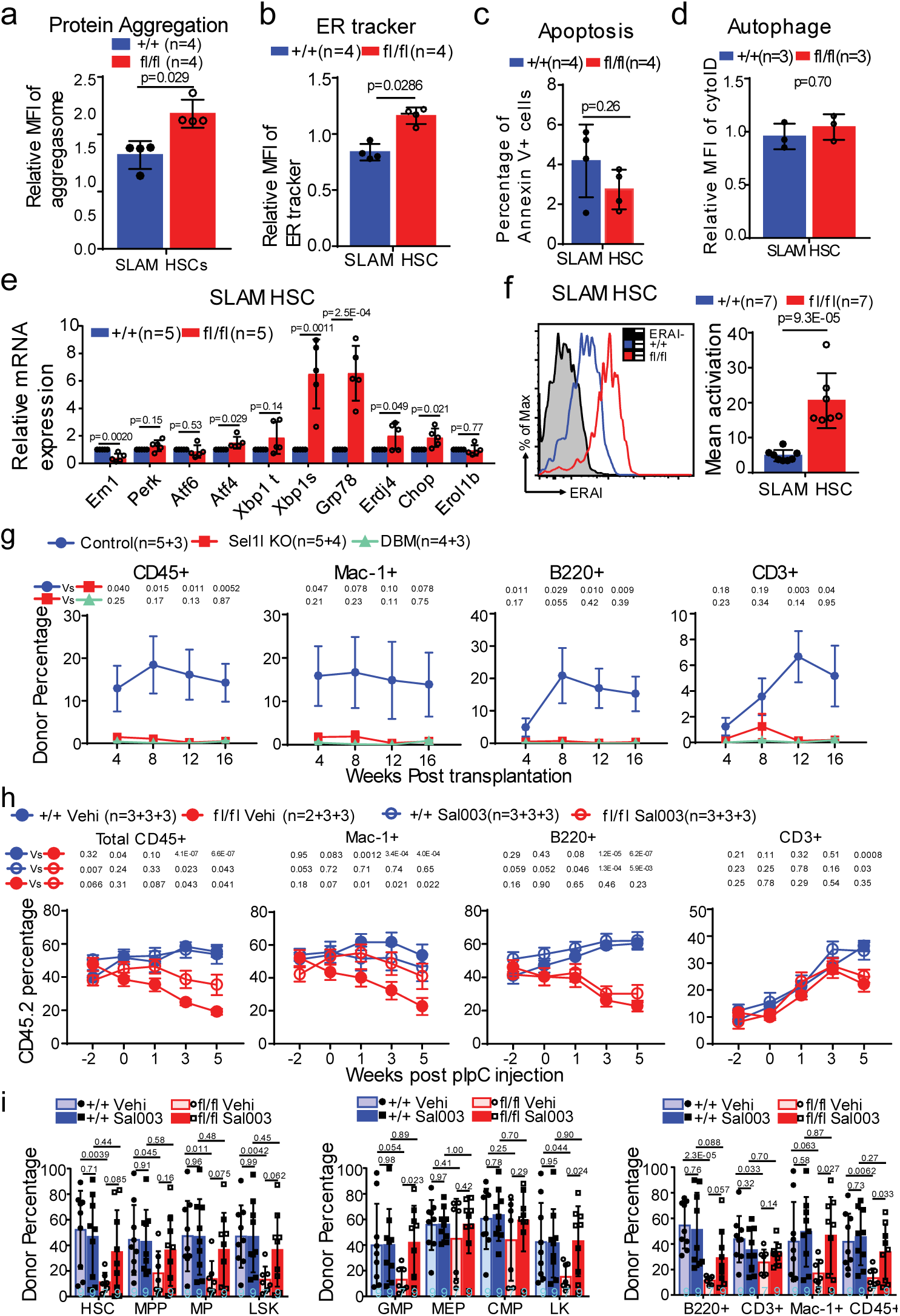
Sel1L-Hrd1 ERAD deficiency leads to ER stress and activation of UPR. **a-e**, Two weeks after pIpC injection, *Mx1-cre^+^; Sel1L^fl/fl^* (fl/fl) or control (+/+) mice were analyzed for protein aggregates (by PROTESTAT, **a**) and ER volume (by ER-Tracker, **b**), apoptosis (by Annexin-V staining, **c**), autophagy activation (by Cyto-ID, **d**) and levels of ER stress signaling targets in purified SLAM HSCs (by RT-qPCR, **e**). **f**, Two weeks after pIpC injection, ERAI *in* SLAM HSCs of *Mx1-cre^+^; Sel1L^fl/fl^; ERAI^+^* (fl/fl) and control (+/+) mice were detect by FACS. Data represent mean fluorescence intensity relative to *ERAI^-^* mice. **g**, Two weeks after pIpC injection, 15 SLAM HSCs purified from control, *Mx1-cre^+^; Sel1L^fl/fl^* (Sel1l KO) and *Mx1-cre^+^; Sel1L^fl/fl^; Ern1^fl/fl^*(DBM) were transplanted with 0.2 million CD45.1 whole bone marrow competitors. The contribution of CD45.2 cells in total CD45+, myeloid (Mac-1^+^), B (B220^+^) and T (CD3^+^) cells was analyzed from peripheral blood. **h-i**, chimerism maintenance analysis reveals rapamycin treatment rescued the loss of reconstitution potential of Sel1L KO HSCs. 5×10^5^ CD45.2^+^ whole bone marrow cells from *Mx1-cre^+^; Sel1L^fl/fl^* (fl/fl) or control mice (+/+) (without pIpC) were transplanted together with 5×10^5^ CD45.1^+^ wild type bone marrow cells into lethally irradiated CD45.1^+^ wild-type recipients. Transplants were injected with pIpC 6 weeks after transplantation and injected with sal003 or vehicle daily starting 6 days before pIpC. **h**, The contribution of CD45.2 cells in total CD45+, myeloid (Mac-1^+^), B (B220^+^) and T (CD3^+^) cells was analyzed from peripheral blood. **i**, Donor contribution to HSCs and other hematopoietic populations was analyzed in transplant recipients from **h**. Note that vehicle treated +/+ or fl/fl data were shared by **h-i** and Figure 6. **b**-**c** as they were performed at the same time to minimize the number of mice used. For **a**-**g**, **i**, data represent mean±s.d., for **h**, data represent mean±sem. For transplant analysis, n is described as numbers of replicates from each round of transplant separated by the plus (+) sign. Two-sided student t-test was used for statistical analysis.

ER stress-induced apoptosis is often associated with increased level of ROS (Han et al., 2013; Zeeshan et al., 2016), which has been shown to impair HSC function and lead to HSC degeneration (Ito et al., 2004; Yahata et al., 2011). Here we found that the level of cellular and mitochondrial ROS remains unchanged in Sel1L KO HSCs (Figure 5i-j). One of the redox regulators, Nrf2, was previously shown to be a substrate of Sel1L/Hrd1 ERAD (Wu et al., 2014), and regulates HSC functions (Tsai et al., 2013). However, the expression levels of key targets of Nrf2, *Hmox*, *Glcm* and *Nqo1*, were not altered in Sel1L KO HSC (**Figure S6c**), suggesting Nrf2 is not involved in regulating ROS level in Sel1L KO HSCs. Autophagy clears the ERAD deficiency-mediated accumulation of protein aggregates (Feng et al., 2017; Quiroga et al., 2013), and was shown to regulate HSC function (Mortensen et al., 2011). We, however, did not detect difference of autophagy by Cyto-ID staining in Sel1L KO HSCs (Figure 6d). Taken together, our results suggest that Sel1L-Hrd1 ERAD deficiency leads to a non-apoptotic ER stress and induces depletion of HSCs independent of ROS or autophagy.

To determine if UPR signalings are activated in Sel1L KO HSCs, we examined the key targets of UPR pathways in purified Sel1L KO. The expression levels of Grp78, total Xbp1, Xbp1s, Erdj4 and Chop were significantly elevated in Sel1L KO HSCs as compared to control HSCs (Figure 6e), indicating that Sel1L loss triggers ER stress and activates UPR. Furthermore, we observed increased phosphorylation of eIF2α, indicative of activated PERK, and increased level of cleaved and activated form of ATF6 (ATF6c) in Sel1L KO LSKs (Figure 7a), suggesting that all three branches of UPR were activated in Sel1L KO HSCs.

**Figure 7.**
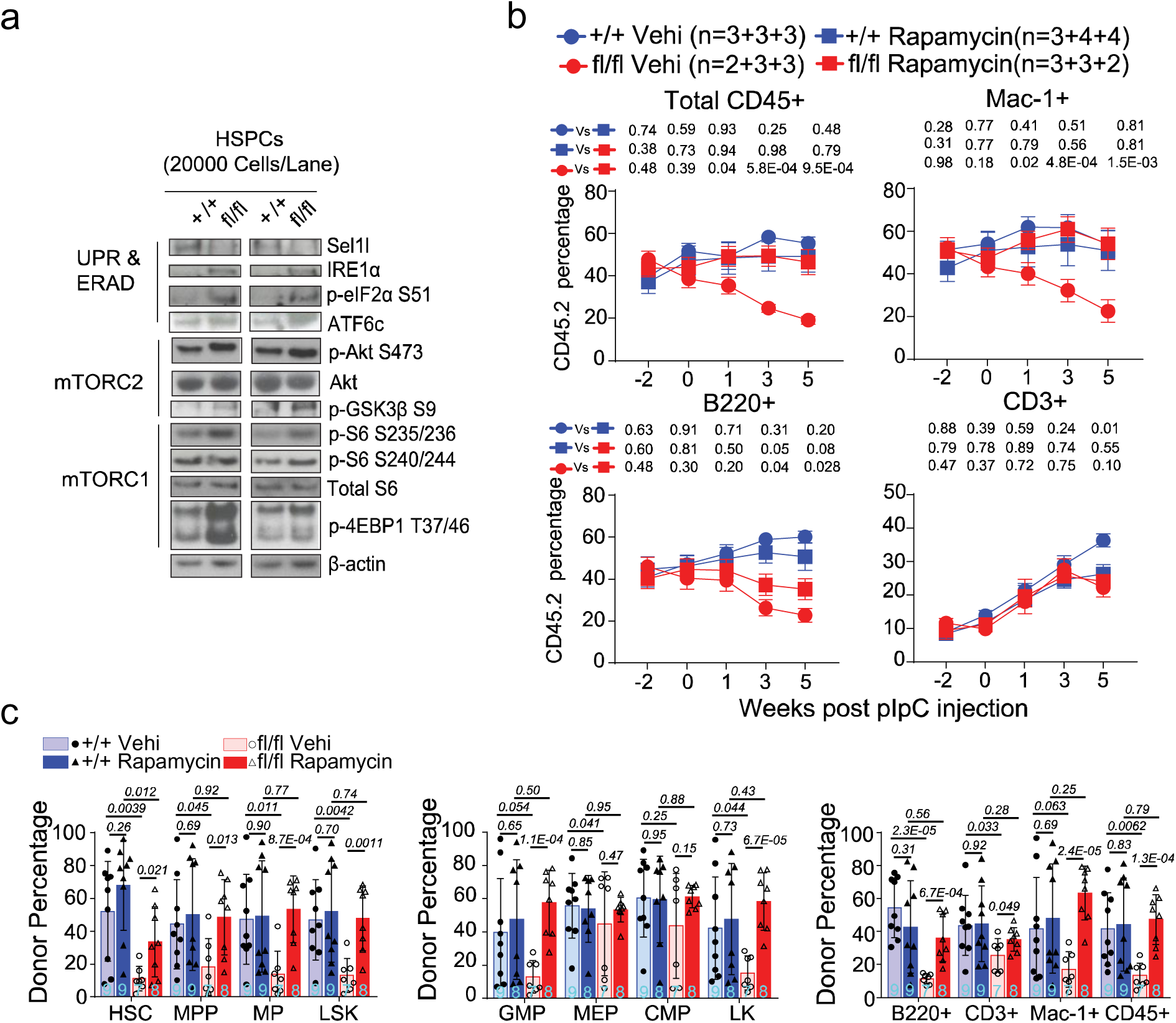
mTORC signaling is activated in Sel1L KO HSCs and inhibition of mTOR rescues HSC reconstitution defects of Sel1L KO. **a**, Western blot of ERAD, UPR, mTORC1/2 proteins in HSPC (LSK). Lysates from equal number of cells were loaded for each lane. **b-c**, chimerism maintenance analysis reveals rapamycin treatment rescued the loss of reconstitution potential of Sel1L KO HSCs. 5×10^5^ CD45.2^+^ whole bone marrow cells from *Mx1-cre^+^; Sel1L^fl/fl^* (fl/fl) or control mice (+/+) (without pIpC) were transplanted together with 5×10^5^ CD45.1^+^ wild type bone marrow cells into lethally irradiated CD45.1^+^ wild-type recipients. Transplants were injected with pIpC 6 weeks after transplantation and injected with rapamycin or vehicle daily starting 6 days before pIpC. **b**, The contribution of CD45.2 cells in total CD45+, myeloid (Mac-1^+^), B (B220^+^) and T (CD3^+^) cells was analyzed from peripheral blood. **c**, Donor contribution to HSCs and other hematopoietic populations was analyzed in transplant recipients from **b**. For transplant analysis, n is described as numbers of replicates from each round of transplant separated by the plus (+) sign. Two-sided student t-test was used for statistical analysis unless specified.

### IRE1α preserves HSC reconstitution under ER stress induced by Sel1L KO

The robust induction of Xbp1s, splicing product of activated IRE1α, and one of Xbp1s target genes, Erdj4, in Sel1L KO HSCs (Figure 6e) suggests activation of IRE1α/Xbp1 signaling. To confirm IRE1α activation, ERAI mice was crossed with *Mx1-Cre^+^; Sel1L^fl/fl^* mice. In ERAI mice, IRE1α activity can be directly monitored by fluorescent expression under the control of the Xbp1 splicing signal (Iwawaki et al., 2004). Compared to control HSCs, Sel1L KO HSCs exhibited much higher level of ERAI, indicating higher IRE1α activity in Sel1L KO HSCs (Figure 6f). To determine whether IRE1α activation mediates Sel1L KO-induced HSC defects, we generated *Mx1-Cre^+^; Sel1lL^fl/fl^*; *Ire1al^fl/fl^* (Sel1L/IRE1α double KO) mice. Two weeks after pIpC injection to knockout Sel1L and IRE1α, we transplanted 15 CD45.2 HSCs from *Mx1-Cre^+^; Sel1l^fl/fl^*, *Mx1-Cre^+^; Sel1L^fl/fl^*; *Ire1α^fl/fl^* (DBM), and wild type control mice, together with 0.2 million CD45.1 protector into lethally irradiated CD45.1 recipients (**Figure S6d**). IRE1α KO did not block the overall loss of donor reconstitution induced by Sel1L KO in peripheral blood (Figure 6g) or bone marrow cells (**Figure S6e**), suggesting that Sel1L KO-induced HSCs defect was not mediated by IRE1α.

Next, to assess the role of IRE1α activation in Sel1L KO HSCs, we sorted the ERAI high (high IRE1α activity) versus ERAI low (low IRE1α activity) Sel1L KO HSCs (**Figure S6f**), and grew them in methylcellulose culture media (M3434) for 10 days. ERAI high Sel1L KO HSCs gave rise to higher number of colonies than ERAI low Sel1L KO HSCs (**Figure S6f**), suggesting that IRE1α activation preserves the function of Sel1L KO HSCs. This protection, however, is not sufficient to correct the Sel1L KO-induced HSC defect in long term reconstitution (Figure 6g and **Figure S6e**). IRE1α has been shown to be a direct degradation target of Sel1L/Hrd1 ERAD (Sun et al., 2015). Consistent with this, the total protein level of IRE1α was increased in Sel1L KO HSC and progenitors (Figure 7a). However, instead of reducing the level of XBP1s as would have been predicted from dose reduction of IRE1α, IRE1α heterozygosity in Sel1L KO HSCs triggered a further increase of XBP1s level (**Figure S6g**), suggesting IRE1α inhibition exacerbates ER stress in Sel1L KO HSCs. Thus, IRE1α plays a protective role in Sel1L KO HSCs.

### Phosphorylation of PERK target eIF2α rescues reconstitution defect of Sel1L KO HSCs

We detected higher level of PERK/eIF2α targets, Aft4 and Chop (Figure 6e), and increased S51 phosphorylation of PERK target, eIF2α, in Sel1L KO HSC and progenitors (Figure 7a), suggesting that Sel1L KO-induced ER stress activated PERK/eIF2α pathway in HSCs. To determine the role of PERK in Sel1L KO, we evaluated the effect of Sal003, an eIF2α dephosphorylation inhibitor, on Sel1L KO-induced HSC defects with chimerism maintenance assay (**Figure S7a**). Sal003 preserves phosphorylation of eIF2α and mimics the effect of PERK-eIF2α activation (Boyce et al., 2005). As shown above, pIpC induced deletion of Sel1L led to a reduction of chimerism of the Sel1L KO cells in transplant recipients, and Sal003 treatment, started 6 days before pIpC and continued throughout the experiment, at least partially blocked the HSC defect induced by Sel1L KO (Figure 6h-i). Pre-treatment with Sal003 did not affect Sel1L deletion in the transplant recipients (**Figure S7b-d**). Taken together, Sel1L KO leads to activation of ER stress, and activation of IRE1α/XBP1s and PERK/eIF2α branches of UPR provides protective effect to Sel1L KO HSCs.

### AKT-mTOR signaling is activated in Sel1L KO HSCs and mediates Sel1L KO-induced HSC impairment

The serine/threonine protein kinase mechanistic target of rapamycin (mTOR) plays important role in an array of cellular functions including protein synthesis, lipid metabolism, mitochondrial functions, cell proliferation, survival and growth (Saxton and Sabatini, 2017). mTOR forms two distinct complexes, mTORC1 and mTORC2, that share common subunits but also contain unique subunits such as Raptor (for mTORC1) and Rictor (for mTORC2)(Saxton and Sabatini, 2017). Activated mTORC1 stimulates mRNA translation and ribosome biogenesis by activating S6K and inhibiting 4EBPs, inhibitors of translation initiation. mTORC2 is activated by growth factors and cytokines, and phosphorylates and stimulates Akt (at S473), to induce mTORC1 activation. Other downstream targets of mTORC2 include FOXO family of transcription factors, serum and glucocorticoid-induced protein kinase 1 (SGK1) and glycogen synthase kinase 3 (GSK3)(Saxton and Sabatini, 2017).

Multiple studies showed that activation of PI3K-Akt-mTOR signaling drives HSCs into proliferation cycle and results in HSC depletion (Chen et al., 2008; Kharas et al., 2010; Yilmaz et al., 2006; Zhang et al., 2006). Similar to Sel1L KO (Figure 5i-j), activation of PI3K-Akt-mTOR by PTEN KO, does not cause ROS accumulation, which is often associated with hyperproliferation of the cell. Given the important role of PI3K-Akt-mTOR in proliferation and metabolic activation, and the similar phenotype of Sel1L KO and Akt-mTOR activation in HSCs, we sought to determine if Akt-mTOR signaling is activated in Sel1L KO HSCs. Western blotting on purified hematopoietic stem and progenitor cells (HSPC; LSK) revealed increased levels of phosphor-S6 (S235/236), phosphor-4EBP1 (T37/46), phosphor-Akt (S473) and phosphor-GSK3β (S9), indicating that both mTORC1 and mTORC2 were activated in Sel1L KO HSPCs (Figure 7a).

To determine the role of mTOR in the HSC dysregulation induced by Sel1L, we inhibited mTOR with rapamycin and determined if this blocked the effect of Sel1L KO in HSCs. We transplanted whole bone marrow cells from *Mx1-cre^+^; Sel1L^fl/fl^* or control mice (without pIpC injection), together with CD45.1 wild-type competitors, into lethally irradiated CD45.1^+^ wild-type recipients (**Figure S7a**). Without pIpC, the Sel1L KO and wild-type HSCs gave rise to similar levels of donor reconstitution in recipients (Figure 7b). Sel1L KO mediated by pIpC led to significant reduction of donor chimerism, but injection of rapamycin (4 mg/kg, i.p daily injection), started 6 days prior to pIpC and continued throughout the experiment, rescued the reduction of donor chimerism in peripheral blood cells from Sel1L KO transplants (Figure 7b). Analysis of bone marrow cells confirmed rescue of donor reconstitution in all hematopoietic populations examined (Figure 7c). Thus, mTOR signaling is activated in Sel1L KO HSCs and mediates the Sel1L KO-induced impairment of HSC reconstitution. Again, pre-treatment with rapamycin did not affect the efficiency of *Sel1L* deletion induced by pIpC injection (**Figure S7b-d**).

In summary, we report here that Sel1L/Hrd1 ERAD is highly active in the most dormant subpopulations of long-term HSCs, and maintains HSC quiescence and self-renewal. Loss of Sel1L drives HSCs into proliferation and activation, accompanied by an increased HSC differentiation and expansion of lineage restricted progenitor populations. Sel1L knockout leads to non-apoptotic ER stress in HSCs, and activation of UPR signaling pathways. Activation of UPR signaling branches IRE1α/Xbp1s and PERK/eIF2α preserves HSC functions upon loss of Sel1L. Finally, mTOR signaling is activated in Sel1L KO HSCs and inhibition of mTOR by rapamycin rescues reconstitution defect of Sel1L KO HSCs. Taken together, protein quality control via Sel1L/Hrd1 ERAD preserves HSC quiescence and self-renewal by restricting mTOR activity.

## Discussion

The protein quality control pathways in ER have been studied in many mammalian cell types, where there is a persistent demand for protein quality control due to high levels of protein synthesis and secretion (Hetz, 2012). In highly specialized cell types, like stem cells that are mostly quiescent and metabolically inactive, the function of protein quality control system remains elusive. Our recent studies, together with other, indicate that protein quality control is critical to preserve the function of HSCs under stress conditions (Liu et al., 2019; van Galen et al., 2014). Here, our studies provide further evidence that protein quality control is required for HSC self-renewal and stemness under steady state when the majority of HSCs are quiescent. Furthermore, Sel1L/Hrd1 ERAD is more active in the most dormant subpopulation of HSCs and maintains HSC in quiescence by restricting mTOR activity. The expression of Sel1L/Hrd1 genes decreases upon HSC proliferation and differentiation, and ERAD deficiency via Sel1L KO leads to profound increase of HSC proliferation, activation and differentiation. Sel1L/Hrd1 ERAD, thus, is actively involved in fate determination of HSCs.

Studies comparing the quiescent and activated HSCs revealed that activated HSCs display much higher levels of expression of proteasome degradation pathway genes (Cabezas-Wallscheid et al., 2017). We show here that although the proteasome degradation pathways are more active in activated HSCs, another branch of protein quality control, ERAD, is much more active in the quiescent subpopulation of HSCs as compared to proliferative HSCs and maintains HSCs in quiescence. Interestingly, a recent study reports that whereas activated neural stem cells (NSCs) have active proteasomes, quiescent NSCs contain large lysosomes to clear protein aggregates from ER to preserve NSC quiescence and functions (Leeman et al., 2018). Furthermore, muscle stem cells exhibit increased level of eIF2α phosphorylation which preserves the self-renewal function of muscle stem cells in vivo (Zismanov et al., 2016). Taken together, protein quality control is critical to maintain stem cell self-renewal and stemness, and different mechanisms are deployed in different types of tissue stem cells to achieve protein homeostasis.

The function of Sel1L/Hrd1 ERAD has been studied in many differentiated cell types including liver, adipocytes and B cells (Bhattacharya et al., 2018; Francisco et al., 2010; Ji et al., 2016; Sha et al., 2014; Sun et al., 2016; Sun et al., 2014; Sun et al., 2015). These studies demonstrated that loss of Sel1L causes dysregulated protein levels of ERAD targets, often membrane associated proteins, which lead to functional impairment of the specific cell type. In many of these cell types, Sel1L loss leads to minimal increase of ER stress. Here we report that in the profoundly quiescent hematopoietic stem cells, Sel1L knockout triggers an overall increase in ER stress and activation of all branches of UPR signaling. Our studies here show that in contrary to many other cell types, Sel1L KO in HSCs activates IRE1α/XBP1s and PERK/eIF2α branches of UPR to preserve HSC functions. Our studies also show that different from other exogenous stress stimulation, such as triggered by tunicamycin or thapsigargin or Lipopolysaccharide (LPS) which induce apoptosis in HSCs (Liu et al., 2019), the ER stress induced by Sel1L KO leads to a non-apoptotic ER stress and represents a physiological stress signal in HSCs.

Our studies identify a previously unknown crosstalk between mTOR signaling and ERAD, and provide new insights into the interactions of cell signaling and protein quality control systems. The role of mTORC1/2 signaling has been studied in apoptotic ER stress and remains controversial (Bobrovnikova-Marjon et al., 2012; Kato et al., 2012; Rajesh et al., 2015). Our data here show that Sel1L KO-induced ER stress activates and depends on mTOR signaling to dysregulate HSCs. The mechanism by which mTOR is activated remains to be investigated. One possibility is that PERK, which has been suggested to activate Akt-mTOR in response to apoptotic ER stress. For instance, PERK/GCN2 mediated eIF2α phosphorylation induces activation of Akt and promotes cell survival under oxidative stress (Rajesh et al., 2015). In addition, PERK was reported to activate Akt-mTOR to promote adipocyte differentiation through a p85 PI3K dependent but eIF2α independent mechanism (Bobrovnikova-Marjon et al., 2012). These raise the possibility that Sel1L knockout-induced mTOR activation is mediated by PERK signaling. In addition, a previous study suggested that mTORC1 and mTORC2 are differentially required for the effect of PTEN knockout in HSCs (Kalaitzidis et al., 2012). Although the main target of rapamycin is mTORC1, prolonged treatment has been shown to inhibit mTORC2 (Sarbassov et al., 2006), and we show here that both mTORC1 and mTORC2 are activated in Sel1L KO HSCs. Future investigations will focus on dissecting the interactions between Sel1L/Hrd1 ERAD, UPR signaling branches and mTOR signaling pathways in HSCs.

## Acknowledgments

This work was supported by the University of Michigan Protein Folding Disease Initiative. Q.L. was supported by NIH/NHLBI (1R01HL132392), American Cancer Society (125080-RSG-13-253-01-LIB), the V Foundation for Cancer Research, Gabrielle’s Angel Foundation, and Leukemia Research Foundation. Thanks to Drs. Hanno Hock, and Masayuki Miura for generously providing *Col1α1-H2B-GFP; Rosa26-M2-rtTA*, and ERAI mice.

## Author Contributions

L.L. performed most of the experiments. A.I., K.F., X.J., M.Z., and G.N. performed some of the experiments with help from L.L. and Q.L. G.S., S.S., and L.Q. contributed to the Sel1L knockout mouse design and analysis. F.M., and Y.D. contributed to analysis of RNAseq database. K.I. contributed to the experiments with mTOR activity and rapamycin treatment. L.L and Q.L. conceived the project, designed experiments, interpreted results, and wrote the manuscript.

## STAR⋆Methods

### Contact for Reagent and Resource Sharing

Further information and request for resources and reagents should be directed and will be fulfilled by the Lead Contact, Dr. Qing Li.

### Experimental Model and Subject Details

#### Mice

All mice were housed in the Unit for Laboratory Animal Medicine (ULAM) at the University of Michigan. All procedures of mouse use and care are in compliance with relevant ethical regulations and policies set forth by the University of Michigan Institutional Animal Care and Use Committee (IACUC), and all animal procedures and protocols have been reviewed and approved by IACUC. Animal protocol was initially approved in 2011 and has been renewed every 3 years since the initial approval. All the mice used in this study were listed in **Key Resources Table** and maintained in C57BL/Ka-CD45.2: Thy-1.1 background. Recipients in reconstitution assays were adult C57BL/Ka-CD45.1: Thy-1.2 mice, at least 8 weeks of age at the time of irradiation. pIpC was reconstituted in PBS and administered at 0.5 µg/gram body mass/day by intraperitoneal injection. All the mice were then analyzed at age 8-14 weeks, a minimum of two weeks after pIpC treatment and paired with sex- and age-matched controls. Doxycycline was added to the water at a concentration of 0.2% (m/v) along with 1% sucrose. All analyses were performed on samples from age- and sex-matched mice. We performed analysis separately for male and female mice and did not observe any differences between genders. Results represent analyses using both male and female mice. Equal number of male and female mice were used for analyses as much as possible.

#### Methods Details

##### Flow cytometry and isolation of hematopoietic cells

Bone marrow cells were flushed from the long bones (tibias and femurs) with FACS Buffer (Hank’s buffered salt solution without calcium or magnesium, supplemented with 2% heat-inactivated calf serum) from the mice. Cells were triturated and filtered through a nylon screen to obtain a single-cell suspension. Hematopoietic populations were analyzed using the antibodies cocktails indicated in **Supplementary Table 2** as previously described(Kiel et al., 2005a; Kiel et al., 2008). For isolation of HSCs, whole bone marrow cells were incubated with antibodies cocktails indicated in **Supplementary Table 2**. After washing, cells were incubated with anti-APC conjugated to paramagnetic microbeads. The microbead bound (c-kit^+^) cells were then enriched using MS columns (Miltenyi Biotec). Non-viable cells were excluded from sorts and analyses using the viability dye 4′,6-diamidino-2-phenylindole (DAPI) (1 μg/ml). Cells were analyzed with BD Fortessa or sorted by BD Aria III flow cytometry. Additional information of antibodies is provided in **Key Resources Table**.

##### Genotyping

150 uL 50 mM NaOH was added into a 1.5 ml tube containing 1-2 mm portion of tail or at least 1 million cells of each mouse. Each sample was heated to 90 °C for 1 hours. 50 uL of Tris-Hcl (pH=8) was then added to each tube to neutralize the reaction. All the samples were then vortexed and 2 ul of each sample was for each genotyping PCR reaction. GoTaq® Green Master Mix was used for Genotyping PCR following the instructions. All the primers were listed in **Supplementary Table 1.**

##### Competitive repopulation assay

Adult recipient mice (CD45.1) were irradiated with an Orthovoltage X-ray source delivering approximately 300 rad min^−1^ in two equal doses of 550 rad, delivered at least 3 h apart. Cells were injected into the tail veins of anaesthetized recipients. Beginning 4 or 5 weeks after transplantation and continuing for at least 16 weeks, blood was obtained from the tail veins of recipient mice, subjected to ammonium-chloride potassium red cell lysis buffer (8 g/L NH4Ac, 1 g/L KHCO3, 0.04 g/L EDTA), and stained with directly conjugated antibodies to CD45.2 (104), CD45.1 (A20), B220 (6B2), Mac-1 (M1/70), CD3 (KT31.1) and Gr-1 (8C5) to monitor engraftment (Lineage Bleeding Cocktail in **Supplementary Table 2**). The chimerism of stem cells, progenitors and lineage cells were analyzed at lead 16 weeks after transplantation unless indicated. Additional information of antibodies is provided in **Key Resources Table**.

##### Chimerism maintenance assay with inhibitors treatment

Adult recipient mice (CD45.1) were irradiated as describe above. One million 1:1 ratio of donor (CD45.2, un-pIpCed) and recipient (CD45.1) Cells were mixed and injected into the tail veins of anaesthetized recipients. Five weeks after transplantation, transplanted recipient mice were injected with vehicle or inhibitors daily along the whole experiments. 6 weeks after transplantation (one weeks after inhibitor injections started), 6 doses of pIpC were injected into all the recipient mice. All the mice were taken down for bone marrow chimerism analysis 6 weeks after inhibitor injections started.

##### Western blotting

The same number of cells (20,000 cells ^-^LSK cells) from each mouse to be analyzed were sorted directly into SF-03 medium without cytokines. The cells were then washed once with SF-03 medium and incubated with 10 ng/ml TPO and 10 ng/ml SCF at 37°C for 10 minutes. The cells were then washed with PBS and precipitated with trichloroacetic acid (TCA) at a final concentration of 10% TCA. Extracts were incubated on ice overnight and spun down for 10 min at 16,000 *g* at 4 °C. The supernatant was removed and the precipitated pellets were washed with ice-cold acetone twice then air-dried. The proteins in the pellets were solubilized with solubilization buffer (9 M Urea, 2% Triton X-100, 1% DTT) before NuPAGE™ LDS sample buffer was added. Proteins were separated on a NuPAGE™ 4-12% Bis-Tris Protein Gels and transferred to a PVDF Membranes. The antibodies used were list in **Supplementary Table 1.**

##### Protein aggregation Assay

Protein aggregation assay was performed by collecting 20 million whole bone marrow cells and staining them with FACS antibodies against cell surface markers for HSCs and other hematopoietic populations as described in **Supplementary Table 2**. After washing, cells were resuspended and fixed with Cytofix/Cytoperm™ Buffer (from BD Pharmingen™ BrdU Flow Kits) on ice for 10 minutes. Cells were then permeabilized with BD Cytoperm™ Permeabilization Buffer Plus (from BD Pharmingen™ BrdU Flow Kits). After washing with 1X Perm/Wash™ Buffer (from BD Pharmingen™ BrdU Flow Kits), the cells were resuspended with 1X Perm/Wash™ Buffer containing 1/5000 PROTEOSTAT^®^ detection reagent and stained for 30 minutes. After washing with FACS Buffer, cells were then resuspended in FACS Buffer containing 2.5 μg/ml DAPI. Aggregation was detected with channel 582/15(488) by BD Fortessa flow cytometry.

##### *In vivo* and *In vitro* Annexin V staining of HSCs

PE Annexin V Apoptosis Detection Kit I was used for Annexin V staining *in vivo*. For *in vivo* staining, 20 million whole bone marrow cells were collected and stained with the cocktail of FACS antibodies as described in **Supplementary Table 2**. After washing with FACS buffer, cells were washed with 1X Annexin V buffer and resuspended in 200 μl 1X Annexin V buffer. PE-conjugated anti-Annexin V antibody was added at a concentration of 1:40 along with 2.5 μg/ml DAPI. For *in vitro* staining, at least 1000 HSCs or CD48-LSK cells were sorted into SF-03 media contained 100 ng/ml TPO/SCF. Cells were recovered overnight before treated with tunicamycin or thapsigargin for 18 hours. Cells were then stained with APC-Annexin V (1:40) along with 2.5 μg/ml DAPI.

##### CFU assay

For colony formation assays, 200 SLAM HSCs were directly sorted into MethoCult^®^ GF M3434 media (Stem Cell technologies, 3434) supplemented with 100 ul SF-03 media containing 100 ng/ml TPO on the top. Cells were then mixed thoroughly by vortex and were plated onto culture dish to grow for 7 days before colonies were counted.

##### RT-qPCR

HSCs were FACS-sorted directly into 500 ul TRIzol™ Reagent, RNA exaction was performed following the instructions. 12 ul Linear Acrylamide was used to help precipitate RNA. Exacted RNA was dissolved in DPEC water and then reverse-transcripted to cDNA following the instruction of High-Capacity RNA-to-cDNA™ Kit. The cDNA was then used for qPCR using Power SYBR™ Green PCR Master Mix. All the qPCR primers were list in **Supplementary Table 1.**

##### Neutral lipid, ER tracker, Mitochondrial staining

20 million whole bone marrow cells and staining them with FACS antibodies against cell surface markers for HSCs and other hematopoietic populations as described in **Supplementary Table 2.**

##### Quantification and statistics analysis

Multiple independent experiments were performed to verify the reproducibility of all experimental findings. All quantitative data represents mean±standard deviation, unless stated otherwise. Unless otherwise indicated, two-tailed Student’s t-tests were used to assess statistical significance for two group comparisons and ANOVA test were performed for comparisons of more than 2 groups, using Prism 7 (GraphPad software). No randomization or blinding was used in any experiments. Experimental mice were not excluded in any experiments. In the case of measurements in which variation among experiments tends to be low (e.g. HSC frequency) we generally examined 3-13 mice. In the case of measurements in which variation among experiments tends to be higher (e.g. reconstitution assays) we examined larger numbers of mice (5-20). Sample size defined as number of independent experiments was described in figure legends. For transplantation experiments, 2-3 donor cells from each genotype were transplanted into 3-5 CD45.1 recipients each donor, and all replicates were pooled for the analysis.

## Data availability

All other data supporting the findings of this study are available from the corresponding author upon reasonable request.

## Key Resources Table

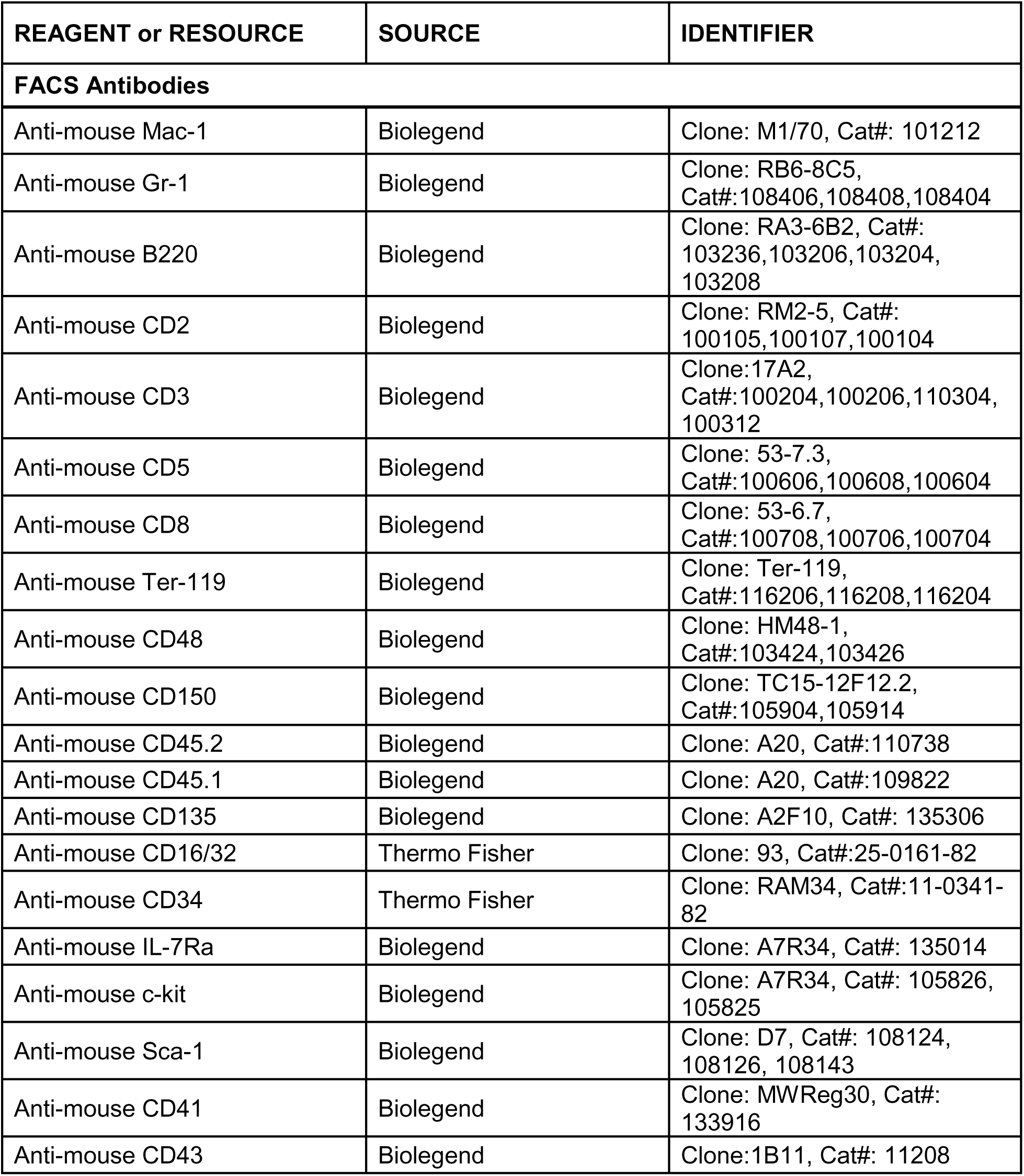
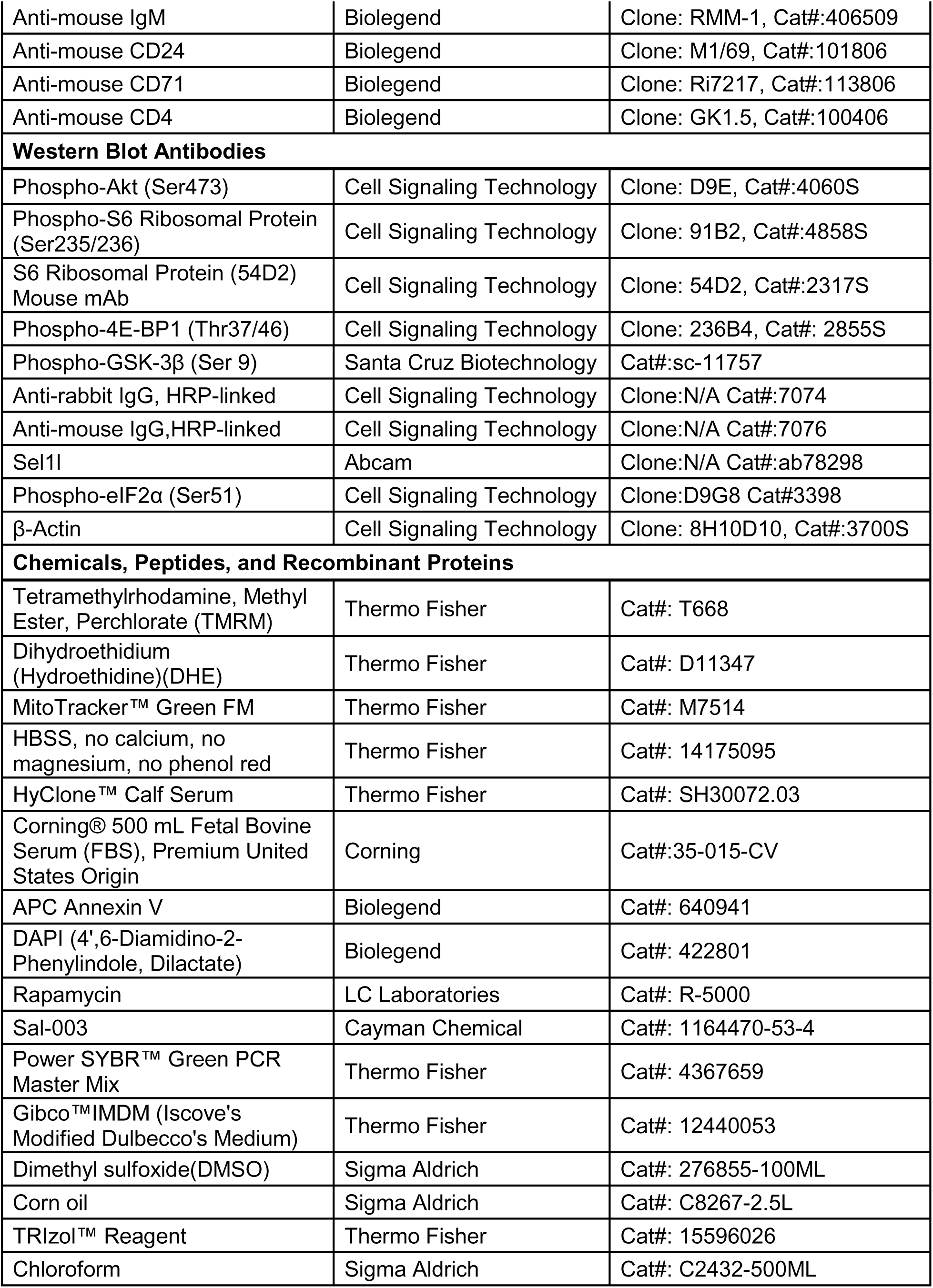
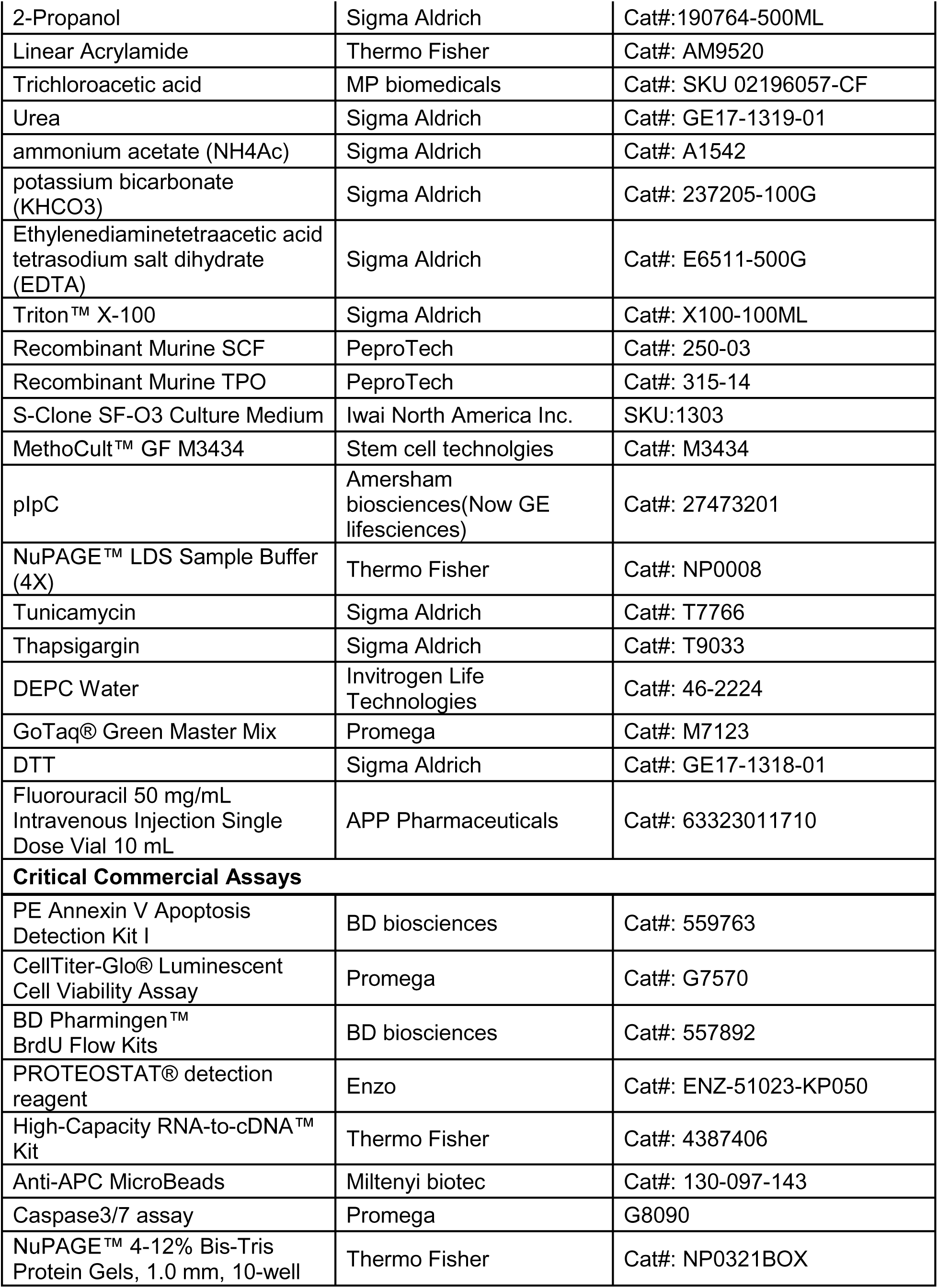
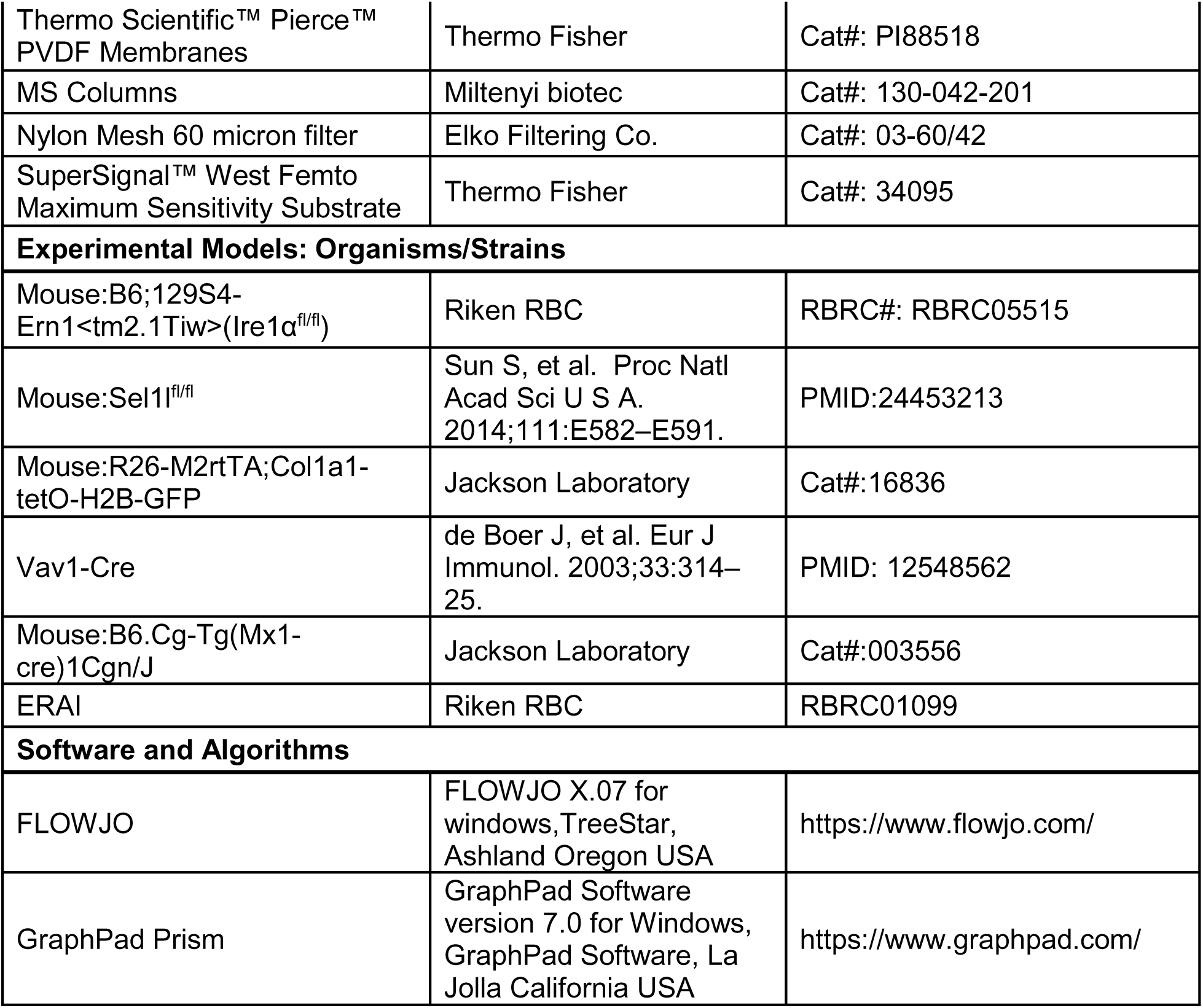

