## supplementary information for "Endoplasmic reticulum associated degradation preserves hematopoietic stem cell quiescence and self-renewal by restricting mTOR activity"

**Supplemental Information**

**Figure S1. Expression of ERAD genes in hematopoietic populations.**

**a,** Representative scheme of doxycycline "labeling-off chase" experiments involved in this study. *Mx-cre*$^{+}$*; Sel1l*$^{fl/fl}$*; Col1α1-H2B-GFP*$^{+/+}$*; Rosa26-M2-rtTA*$^{+/+}$ *or Mx-cre*$^{-}$*; Sel1l*$^{fl/fl}$*; Col1α1-H2B-GFP*$^{+/+}$*; Rosa26-M2-rtTA*$^{+/+}$ *mice* were pIpC injected, doxycycline was added to drinking water at least 2 weeks after pIpC injection. Note that *Mx-cre*$^{-}$*; Col1α1-H2B-GFP*$^{+/-}$*; Rosa26-M2-rtTA*$^{+/-}$ mice were used for **Figure 1a-b**, and *Mx-cre*$^{-}$*; Col1α1-H2B-GFP*$^{+/+}$*; Rosa26-M2-rtTA*$^{+/+}$ were used for **Figure 1C**. **b**, Re-analysis of ERAD gene expression in dHSC, aHSCs and MPP1 on previously published RNAseq database (Cabezas-Wallscheid et al., 2017, ArrayExpress: E-MTAB-4549). **c-d**, Representative FACS plots of TMRM staining of whole bone marrow cells from 24 weeks "off label chase" *of Col1α1-H2B-GFP*$^{+/+}$*; Rosa26-M2-rtTA*$^{+/+}$ mice. **e**, RT-qPCR of ERAD genes in different populations of 8-10 weeks old wild type mice. Data represent mean±s.d. Two-sided student t-test was used for statistical analysis.

**Figure S2. Steady state hematopoiesis in *Mx-cre*$^{+}$*; Sel1l*$^{fl/fl}$ mice (related to Figure 2).**

Two weeks after pIpC injection, *Mx-cre*$^{+}$*; Sel1l*$^{fl/+}$(fl/+), *Mx-cre*$^{+}$*; Sel1l*$^{fl/fl}$(fl/fl), and *Mx-cre*$^{-}$*; Sel1l*$^{fl/fl}$(+/+) were analyzed. **a,** Expression of Sel1l in whole bone marrow cells. **b-d,** Representative body, spleen and thymus photo of *Mx-cre*$^{+}$*; Sel1l*$^{fl/fl}$(fl/fl), and *Mx-cre*$^{-}$*; Sel1l*$^{fl/fl}$(+/+). **e-m,** Hematopoietic cell populations frequency in bone marrow, spleen and thymus of *Mx-cre*$^{+}$*; Sel1l*$^{fl/fl}$(fl/fl), and *Mx-cre*$^{-}$*; Sel1l*$^{fl/fl}$(+/+).Data represent mean±s.d. Two-sided student t-test was used for statistical analysis.

**Figure S3. Steady state hematopoiesis in *Vav-cre*$^{+}$*; Sel1l*$^{fl/fl}$ mice (related to Figure 3)**

6-8 weeks old *Vav-cre*$^{+}$*; Sel1l*$^{fl/fl}$(fl/fl), and *Vav-cre*$^{-}$*; Sel1l*$^{fl/fl}$(+/+) were analyzed. **a-d,** Representative body, spleen and thymus and long legs photo of *Mx-cre*$^{+}$*; Sel1l*$^{fl/fl}$(fl/fl), and *Mx-cre*$^{-}$*; Sel1l*$^{fl/fl}$(+/+). **e-j,** Hematopoietic cell stem cell populations frequency in bone marrow, spleen of *Vav-cre*$^{+}$*; Sel1l*$^{fl/fl}$(fl/fl), and *Vav-cre*$^{-}$*; Sel1l*$^{fl/fl}$(+/+). Data represent mean±s.d. Two-sided student t-test was used for statistical analysis

**Figure S4. Transplantation experiments in Figure 4.**

Schematic representation of transplantation experiments in **Figure 4**.

**Figure S5**. **Loss of Sel1L leads to decrease of HSC quiescence.**

**a**, two weeks after pIpC injection, SLAM HSCs were purified from *Mx-cre*$^{+}$*; Sel1l*$^{fl/fl}$(fl/fl), and mRNA level of p27 and p57 were detected by RT-qPCR. Data represent mean±s.d. Two-sided student t-test was used for statistical analysis. **b**, Two weeks after pIpC injection, *Mx-cre*$^{+}$*; Sel1l*$^{fl/fl}$(fl/fl), and *Mx-cre*$^{-}$*; Sel1l*$^{fl/fl}$(+/+) were injected with 150 mg/kg 5-FU weekly, data represents survival of injected mice. Kaplan-Meier test was used for statistical analysis.

**Figure S6. Protective role of UPR in Sel1l KO HSCs (related to Figure 6).**

**a-b,** Representative FACs plot **a** and summary **b** of apoptosis and ERAI activation after treatment with tunicamycin or thapsigargin. Two weeks after pIpC injection,1000 HSCs were purified from *Mx-cre*$^{+}$*; Sel1l*$^{fl/+}$ *ERAI*$^{+}$*;* (+/+)*, Mx-cre*$^{+}$*; Sel1l*$^{fl/fl}$*; ERAI*$^{+}$*;* (fl/fl), after recovery overnight, cells were then treated with Tunicamycin (Tm) and Thapasigargin for 18 hours before apoptosis level was measured. **c,** Two weeks after pIpC injection, SLAM HSCs were purified from *Mx-cre*$^{-}$*; Sel1l*$^{fl/fl}$*;Ern1*$^{fl/+}$*or Mx-cre*$^{-}$*; Sel1l*$^{fl/f}$ (+/+), *Mx-cre*$^{+}$*; Sel1l*$^{fl/fl}$ (fl/fl), mRNA was extracted and NRF2 target gene expression was measured by RT-qPCR. Data represent mean±s.d. Two-sided student t-test was used for statistical analysis. **d**, Schematic representation of transplantation experiments in **Figure 6g**. **e**, Donor contribution to HSCs and other hematopoietic populations was analyzed in the bone marrow 16 weeks after transplantation in **Figure 6g**. Data represent mean±s.d. Two-sided student t-test was used for statistical analysis. **f**, Two weeks after pIpC injection, 200 bulk HSCs or $ERAI^{high}$ sub-population and $ERAI^{low}$ sub-population HSCs from *Mx-cre*$^{+}$*; Sel1l*$^{fl/+}$ *ERAI*$^{+}$*;* (+/+), *Mx-cre*$^{+}$*; Sel1l*$^{fl/fl}$*; ERAI*$^{+}$*;* (fl/fl) were plated into semi-solid Meth-cult media for colony assay. Colony numbers were counted after 10 days of culture. Data represent mean±s.d. Two-sided student t-test was used for statistical analysis. **g**, Two weeks after pIpC injection, SLAM HSCs were purified from *Mx-cre*$^{-}$*; Sel1l*$^{fl/fl}$*;Ern1*$^{fl/+}$ *or Mx-cre*$^{-}$*; Sel1l*$^{fl/fl}$*; Ern1*$^{fl/+}$ (Control), *Mx-cre*$^{+}$*; Sel1l*$^{fl/fl}$*,* or *Mx-cre*$^{+}$*; Sel1l*$^{fl/+}$ *Ern1*$^{fl/+}$ mice, and gene expression was measured by RT-qPCR. Data represent mean±s.d. Two-sided student t-test was used for statistical analysis except for **b**, where two-way Anova test was used for group comparison.

**Figure S7. Inhibitors treatment did not affect the efficiency of Sel1l1 deletion by pIpC in chimerism maintenance experiments.**

**a**, Schematic representation of chimerism maintenance experiments in **Figure 6h-i** and **Figure 7b-c**. **b**, Schematic representation of Floxed allele and wild type allele of *Sel1l*. Genotyping-F and Genotyping-R were used to detect Floxed allele and Wild type allele, there are no bands from recombined Floxed allele. **c**, Representative genotyping results of whole bone marrow cells from *Mx-cre*$^{+}$*; Sel1l*$^{fl/+}$, *Mx-cre*$^{+}$*; Sel1l*$^{fl/+}$ mice after 6 does of pIpC injection. **d**, Representative genotyping results of whole bone marrow cells from *Mx-cre*$^{+}$*; Sel1l*$^{fl/fl}$ (CD45.2), transplanted recipients, after pIpC injection and inhibitors treatment. Loss of the mutant band (286) indicates recombination of the mutant allele and deletion of Sel1L.

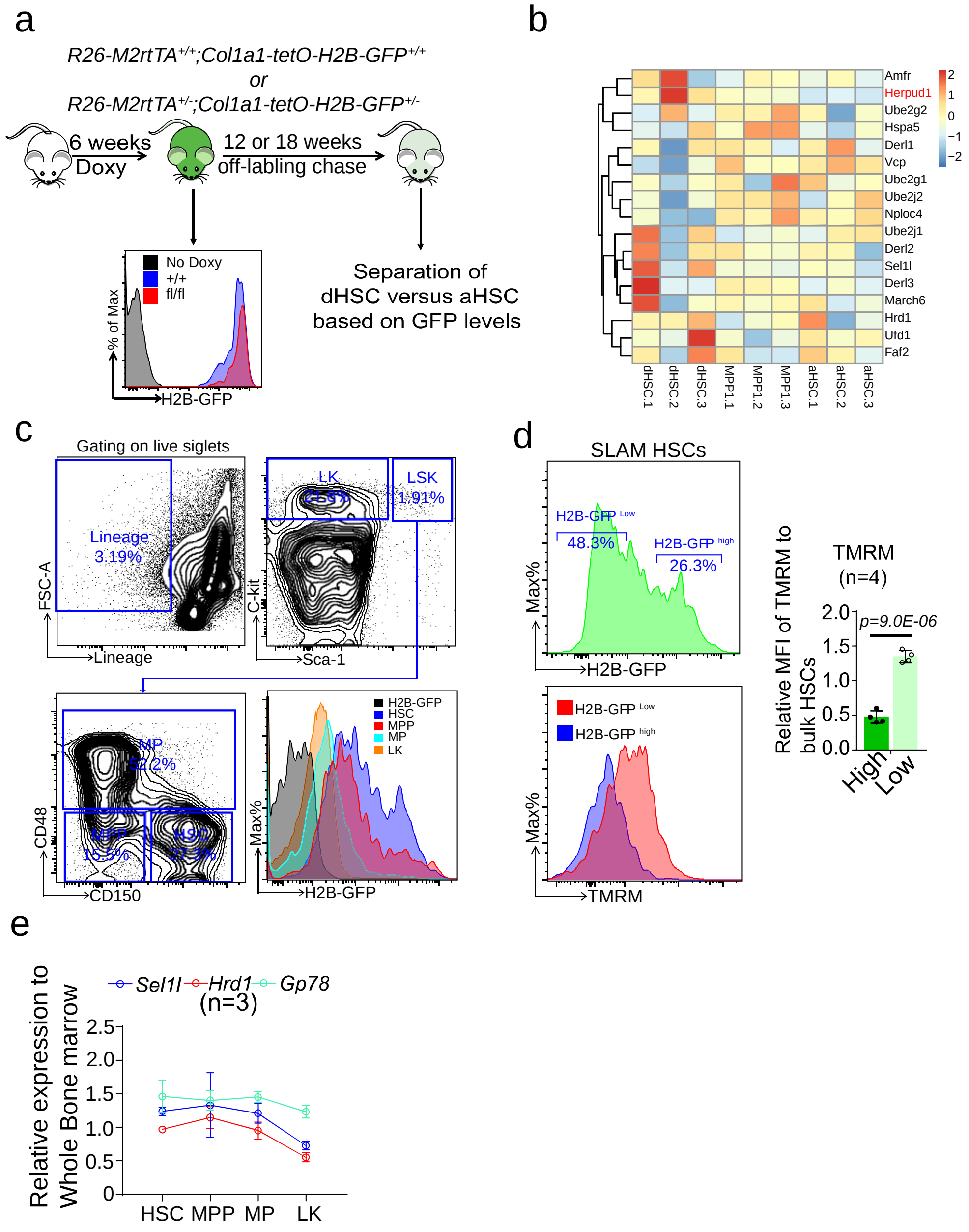


**Figure S1. Expression of ERAD genes in hematopoietic populations (related to Figure 1).**

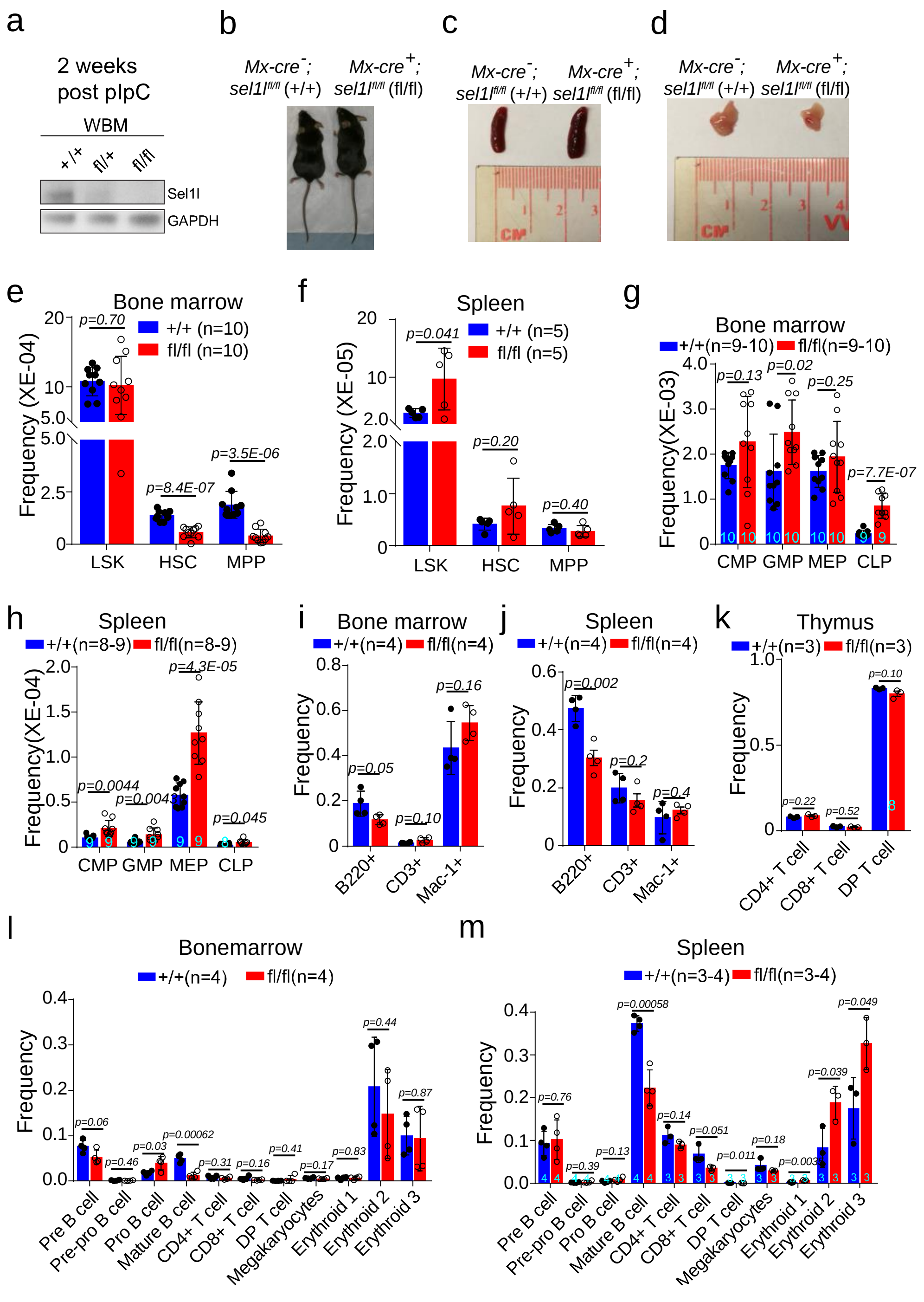


**Figure S2. Steady state hematopoiesis in *Mx-cre*$^{+}$*; Sel1l*$^{fl/fl}$ mice (related to Figure 2).**

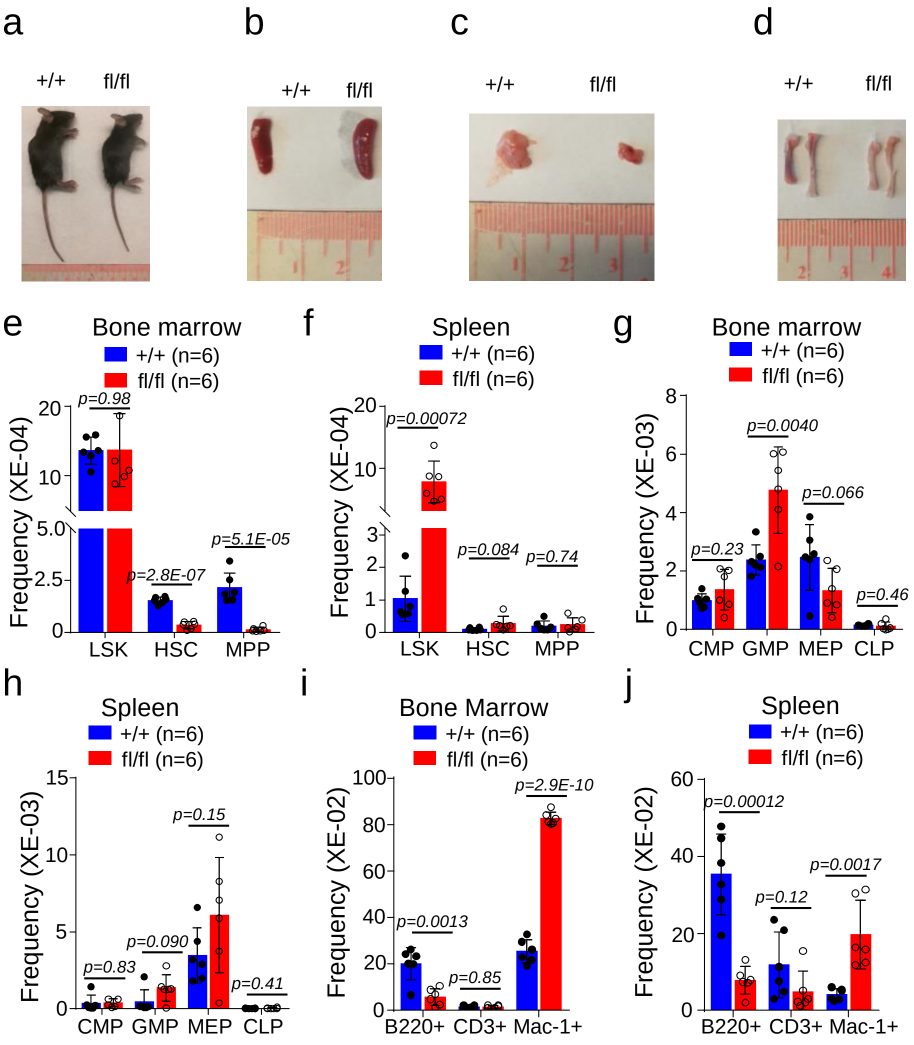


**Figure S3. Steady state hematopoiesis in *Vav-cre*$^{+}$; *Sel1l*$^{fl/fl}$ mice (related to Figure 3)**

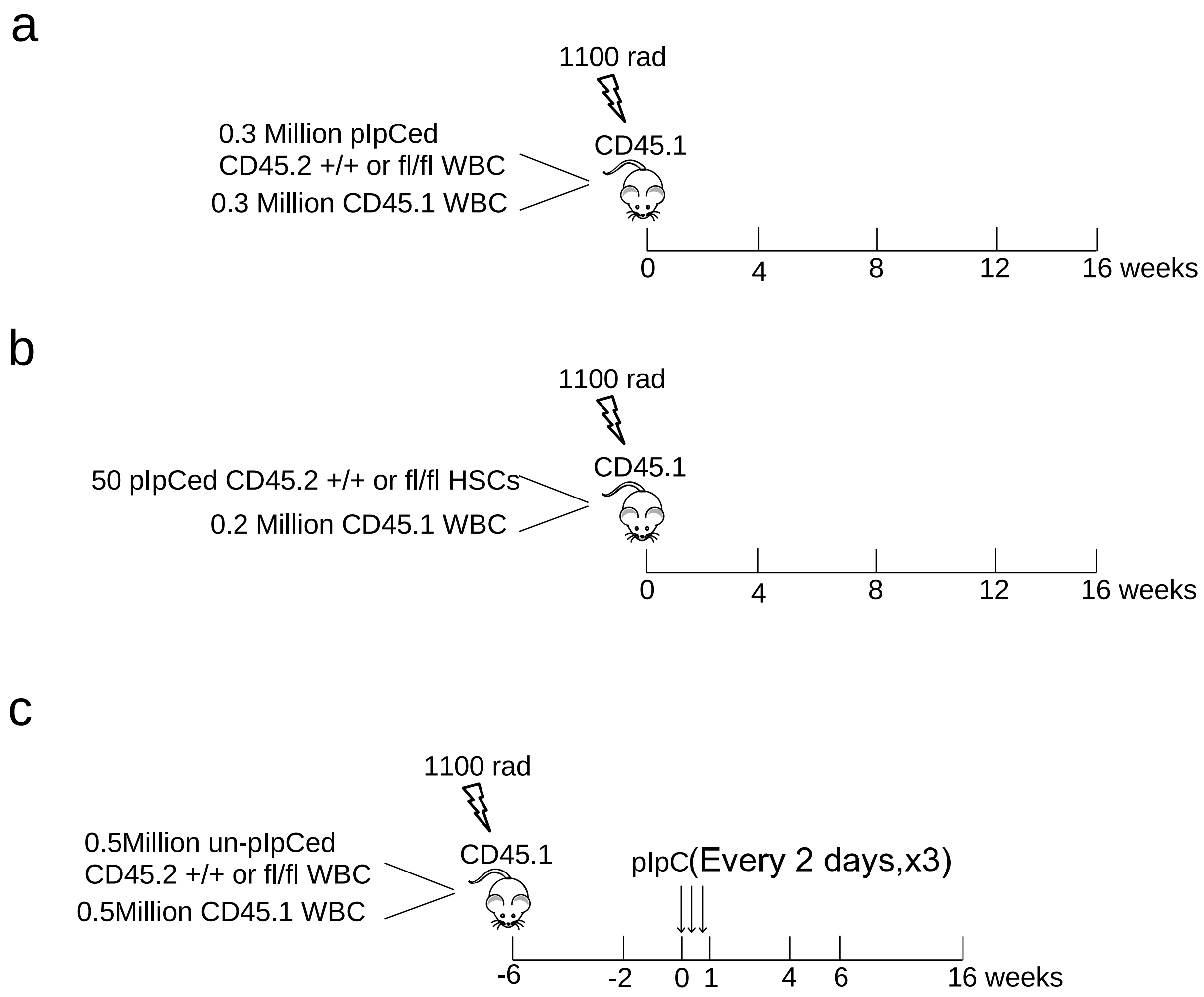


**Figure S4. Transplantation experiments in Figure 4 (related to Figure 4).**

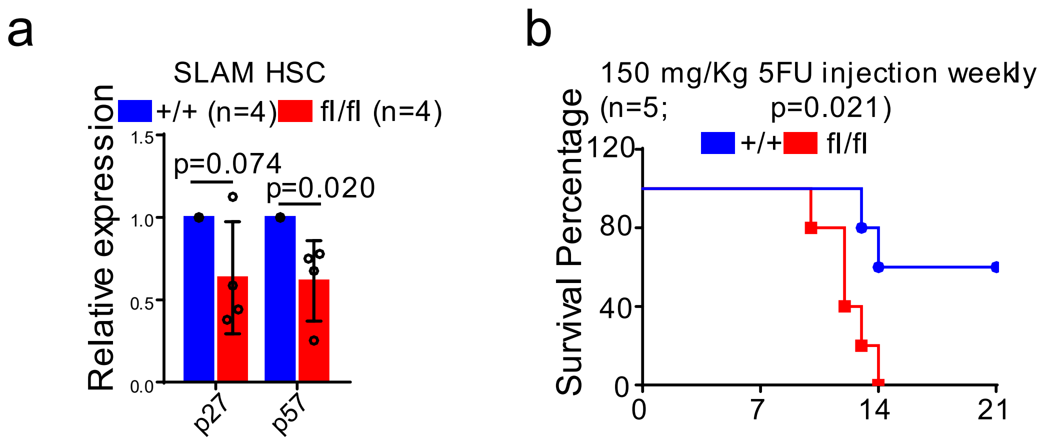


**Figure S5. Loss of Sel1L leads to decrease of HSC quiescence (related to Figure 5).**

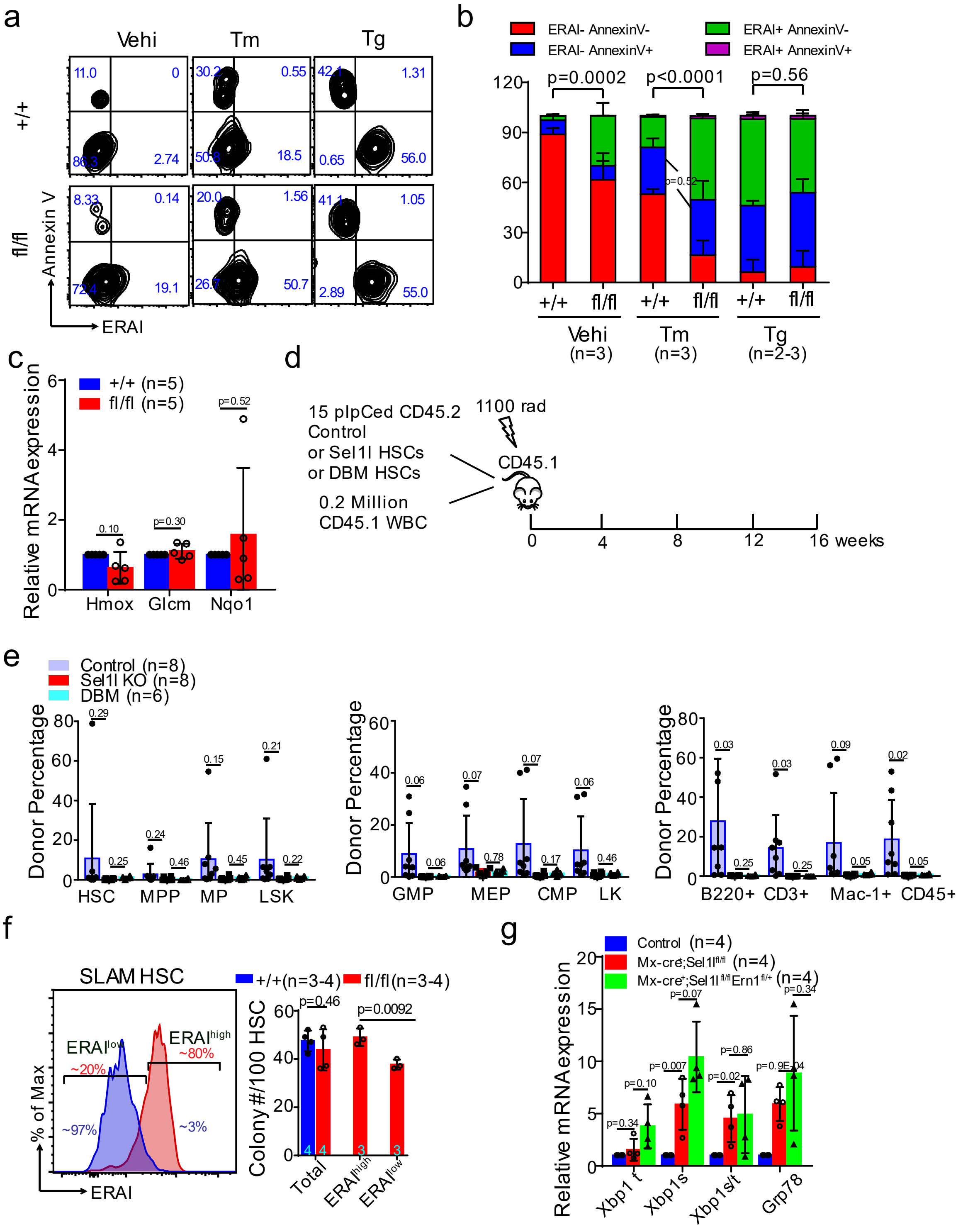


**Figure S6. Protective role of UPR in Sel1l KO HSCs (related to Figure 6).**

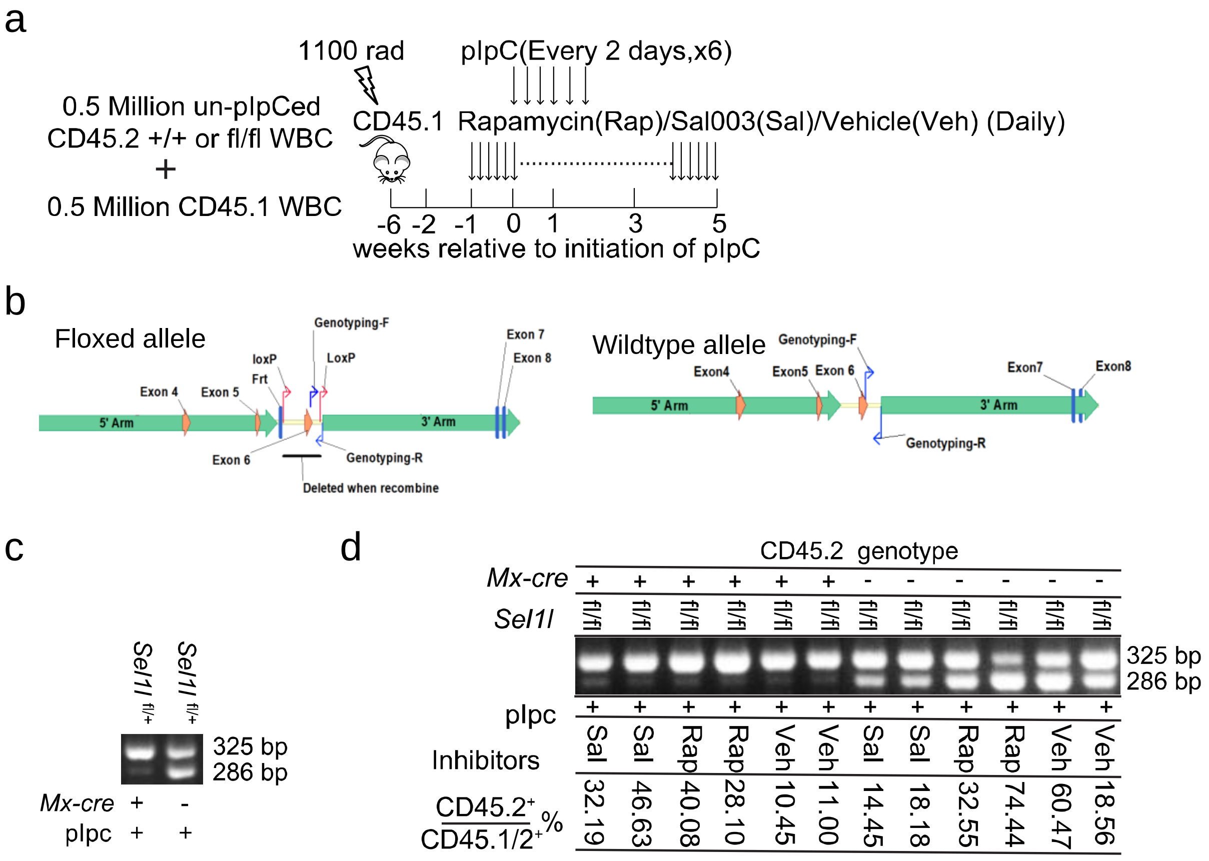


**Figure S7. Inhibitors treatment did not affect the efficiency of Sel1l1 deletion by pIpC in chimerism maintenance experiments.**
